# Assembly of host-specific bacterial and fungal communities from soil by diverse subaerial green algae

**DOI:** 10.64898/2026.11.06.764884

**Authors:** Magdalena W. Slawinska, Barbara Melkonian, Michael Melkonian, José Flores-Uribe, Paloma Durán, Ruben Garrido-Oter

## Abstract

Green algae recruit microbes from their environment to the phycosphere, a region closely surrounding algal cells which is analogous to the rhizosphere of land plants. Recent work showed that subaerial green algae can recruit and assemble their phycosphere microbiota from soil-derived bacteria. However, how widespread this ability is among diverse green algae, whether host identity shapes community composition, and whether soil-borne fungi can be recruited to the phycosphere remained unclear. Here, we show that subaerial green algae spanning Chlorophyta and Streptophyta assemble distinct phycosphere communities from soil-derived bacteria and fungi, with host identity as the primary driver of phycosphere composition, both in natural soil and in a liquid mesocosm. In soil, bacterial communities were strongly reshaped by algal hosts, whereas the fungal communities responded more weakly and less consistently. In liquid system, bacterial community composition was correlated with host phylogeny. Phycosphere communities included taxa known to associate with land plants in the rhizosphere, supporting the finding that some microbial taxa associate with a wide range of photosynthetic hosts. Our findings demonstrate that the green algae capacity to assemble distinct phycosphere communities possibly predates the divergence of Chlorophyta and Streptophyta. The stronger effect of host identity on bacterial than fungal community mirrors patterns observed in land plant microbiota, suggesting that principles of host-microbe interactions are conserved across evolutionarily distant photosynthetic hosts.

## Introduction

In nature, algae recruit microbes from their environment to their phycosphere, a region closely surrounding algal cells (Seymour *et al*., 2017). The phycosphere is an environment analogous to the rhizosphere of land plants, and at both interfaces, exudates alter the physicochemical environment, stimulating bacterial colonization (Seymour *et al*., 2017). Among the colonizing bacteria, some can be found at both interfaces (Seymour *et al*., 2017; Durán *et al*., 2022). Furthermore, in exchange for carbon and metabolites provided by the algal host (Durham *et al*., 2015; Calatrava *et al*., 2024), microbial communities supply their photosynthetic hosts with micronutrients (Croft *et al*., 2005; Amin *et al*., 2009; Castrillo *et al*., 2017; Kimbrel *et al*., 2019; Harbort *et al*., 2020), can alleviate stresses (Xu *et al*., 2018; Simmons *et al*., 2020) or provide protection against pathogens (Durán *et al*., 2018; Carrión *et al*., 2019; Flores *et al*., 2024).

Previously, research on interactions between microscopic algae and their phycosphere communities focused mainly on aquatic environments and the interactions with their associated bacteria (Seymour *et al*., 2017; Kimbrel *et al*., 2019; Fei *et al*., 2020; Kim *et al*., 2022). It has been estimated that in aquatic environments, algae and bacteria engage in metabolic interactions, with bacteria consuming up to 50% of the carbon fixed by algae, and algae profiting from micronutrients provided by bacteria (Croft *et al*., 2005; Amin *et al*., 2009; Amin *et al*., 2015; Durham *et al*., 2015; Paerl *et al*., 2016). Algal photosynthesis is also important in terrestrial environments, where soil algae take up carbon in amounts equal to ca. 31% of the global anthropogenic carbon emissions (Friedlingstein *et al*., 2019; Jassey *et al*., 2022). However, despite their importance for the global carbon cycle and mitigating climate change, and the dependence of many algae on bacterial presence for optimal growth (Amin *et al*., 2009; Jassey *et al*., 2022), the interactions of soil-borne microscopic algae with their associated microbial communities remain underexplored.

In our previous work, we showed that the soil-derived model alga *C. reinhardtii* (Sasso *et al*., 2018) and a small number of other chlorophyte and streptophyte algae can recruit and assemble a phycosphere bacterial community from soil-borne bacteria (Durán *et al*., 2022). These observations with the hypothesis that the common ancestor of land plants and algae was preadapted for symbiosis, and that the ability to associate with microbial communities facilitated the land colonization by plants (Delaux *et al*., 2015; Knack *et al*., 2015; Delaux and Schornack, 2021). However, whether this ability is widespread among subaerial green algae, extends to soil-borne fungi, and if community composition mirrors host phylogeny (phylosymbiosis), remain unknown.

Here, we assembled a taxonomically broad collection of different subaerial green algae, including the model alga *C. reinhardtii* and microscopic algae originating from the same environment, encompassing Chlorophyta (*Chlorophyceae*, *Ulvophyceae*, *Trebouxiophyceae*) and Streptophyta algae (*Klebsormidiophyceae*, *Zygnematophyceae*), with *Spirogloea muscicola* representing the algal lineage believed to be the closest extant relative to the most recent common ancestor of streptophyte algae and land plants (Cheng *et al*., 2019). First, we used this collection to characterize the bacterial and fungal communities associated with the subaerial algae in a terrestrial environment. Second, we explored the temporal dynamics in the bacterial communities in controlled experiments in liquid cultures. Together, we show that host identity is the main driver of phycosphere composition and that the host effect is more pronounced for bacterial than for fungal communities. Furthermore, we show that host-driven differences in bacterial community composition are substantial and consistent between the two experimental setups used. Our findings indicate that the ability to associate with soil-derived bacteria and fungi is widely found among subaerial green algae ranging from Chlorophyta to Streptophyta, and that host identity is the main driver of phycosphere composition.

## Results

### Microbial diversity in the native environment and reference soil

We first aimed to determine whether subaerial algae associate with distinct bacterial and fungal communities in natural soil, and whether the bacterial and fungal community composition is correlated with host phylogeny. To this end, we isolated a subset of algal strains originating from the same terrestrial environment in a natural site located in the Eifel National Park, Germany. This collection includes chlorophyte and streptophyte algae belonging to different classes, capturing a broad phylogenetic diversity (Supplementary Table 1). To address these questions under controlled conditions, we designed experiments with a soil (Cologne Agricultural Soil, CAS) sourced near the Eifel National Park, which we used before for the isolation and characterization of phycosphere bacteria (Durán *et al*., 2022). Because CAS stands in for the algae’s native substrate in these experiments, we first asked whether its microbial pool is representative of the Eifel site.

To compare the bacterial and fungal communities from the native environment of the algae (Eifel site) and our reference CAS soil to assess their similarity we used amplicon sequencing of the V5-V7 16S rRNA gene and internal transcribed spacer (ITS2) regions. As expected, fungal and bacterial communities from the Eifel site showed higher alpha-diversities compared to those from CAS, indicating a larger microbial diversity present in this natural site (**Fig. 1a-b**). Microbial community structures differed between the sites for both bacteria and fungi, with the variable *site* explaining 16.1% and 11.4% of the overall variance of the data, respectively (Constrained Principal Coordinate Analysis, CPCoA, and permutation test, *P* <0.001) (**Fig. 1c-d**). We observed higher heterogeneity among the Eifel samples than among the CAS samples, which can be attributed to the higher complexity of this natural site compared to the uniform, unplanted samples from CAS. To explore which taxa drove the compositional differences between the two sites, we analysed the taxonomic composition at the family level and identified shared taxa, which accounted for 84.2% of the aggregated relative abundance (RA) in the Eifel samples and 80.8% in CAS samples (**Fig. 1e**). Although most bacterial families were found in both sites, their aggregated RA differed in multiple cases (**Fig. 1e**, **Fig. S1**), with Caulobacteraceae and Xanthobacteraceae being less abundant in CAS, and Lysobacteraceae and Pseudomonadaceae being more abundant.

**Figure 1.**
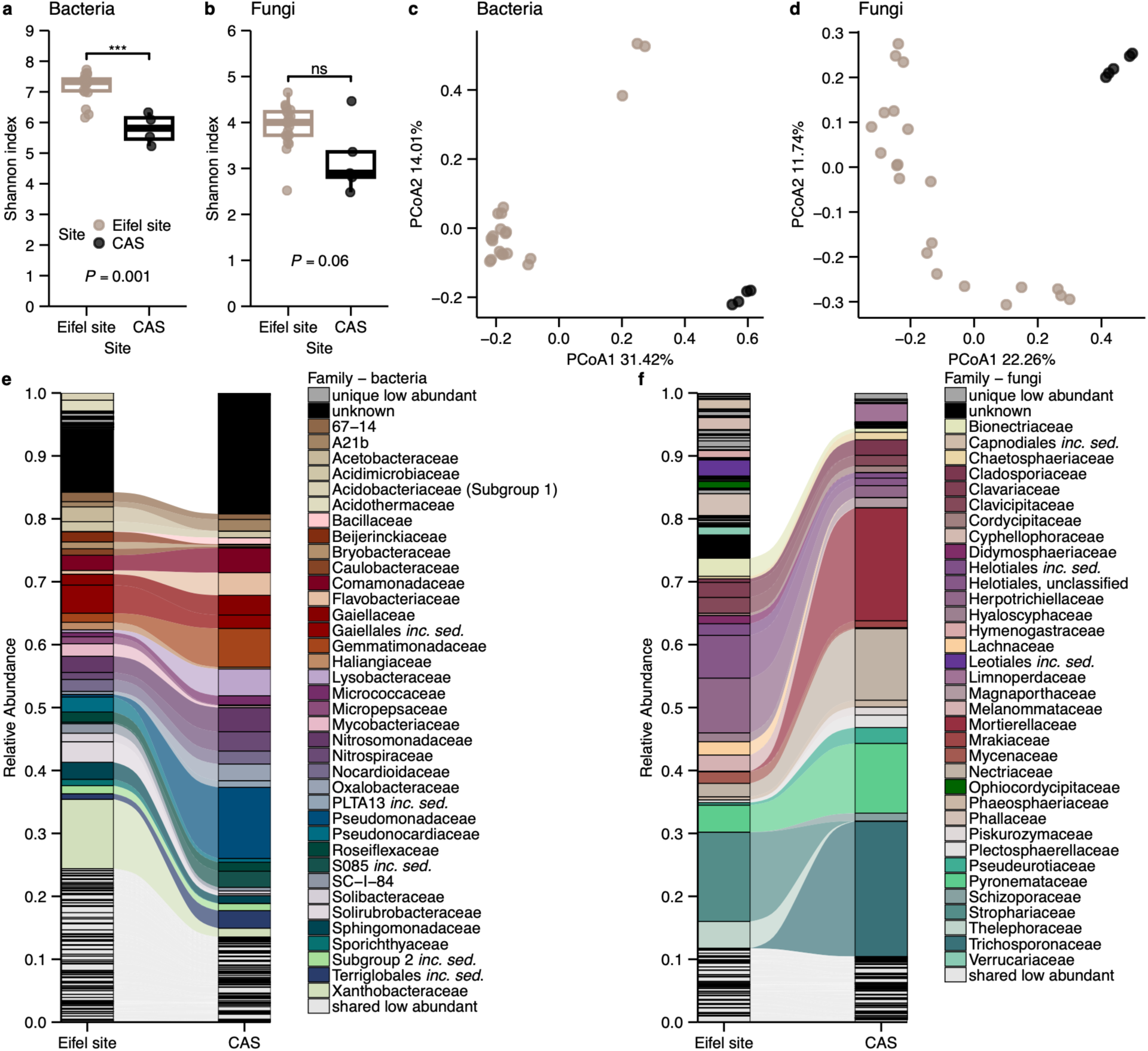
Comparison of bacterial and fungal community structures from the Eifel site and native CAS. (**a**, **b**) Alpha diversity estimates of bacterial (**a**) and fungal (**b**) communities from the Eifel site (*n* = 21 for bacteria and fungi) and CAS (*n* = 4 for bacteria, *n* = 5 for fungi). The two-sided Mann-Whitney test was used to assess significant differences among groups (*P* <0.05). (**c**, **d**) Principal Coordinates Analysis of Bray-Curtis dissimilarities (ASV-level) of bacterial (**c**) and fungal (**d**) communities, with site explaining 16.1% and 11.4 % of variance in community dissimilarity, respectively (constrained PCoA and permutation test, *P* <0.001). The shared legend for panels (**a**-**d**) is presented in panel (**a**). (**e**, **f**) Alluvial plots showing the average relative abundances of bacterial (**e**) and fungal (**f**) families in Eifel site and CAS samples.

Differences in the taxonomic composition between sites were more pronounced in the fungal than in the bacterial communities (**Fig. 1f**, **Fig. S2**). Fungal families shared between the sites accounted for 73.7% aggregated RA in Eifel and 94.4% in CAS, and some of the highly abundant families were unique to one of the sites. Interestingly, the abundances of the dominant fungal families differed markedly between the sites, with the RAs of the three most abundant families in CAS (Mortierellaceae, Pyronemataceae, Trichosporonaceae) adding up to 50.5%, whereas in the Eifel samples, summing up to only 4.4%. These findings indicate that fungal communities associated with subaerial microalgae are more site-specific than bacterial ones, as previously demonstrated for fungal communities associated with land plants (Thiergart *et al*., 2020; Bourceret *et al*., 2022).

Analysis of the bacterial and fungal communities of the Eifel site and CAS showed that the communities of both sites are highly diverse. Although they differ at lower taxonomic levels, they share dominant bacterial phyla and families, indicating that bacterial taxa broadly representative of the algae’s native environment are present in CAS. In contrast, the fungal communities showed greater site-specificity with less compositional overlap, but some similarities between the two sites could be found, supporting the use of CAS as the source of soil microbes for the experiments that follow.

### The model alga *C. reinhardtii* shapes bacterial and fungal communities in a natural soil

We previously observed the ability of the model alga *C. reinhardtii* to associate with soil-derived bacteria, showing parallels to the root microbiota assembly in land plants (Durán *et al*., 2022). To examine whether *C. reinhardtii* also influences soil fungal communities, we inoculated CAS-filled pots with axenic alga cultures and collected samples after the alga had substantially grown on the soil surface (**Materials and Methods**). We then characterized both bacterial and fungal phycosphere microbiomes by amplicon sequencing of the V5-V716S rRNA gene and ITS2 regions. The *C. reinhardtii* phycosphere bacterial community clustered separately from both input CAS samples and the negative controls (soil without *C. reinhardtii* inoculum) (**Fig. 2a**), recapitulating our previous findings (Durán *et al*., 2022). Similarly, fungal community profiles showed significant separation between the *C. reinhardtii* phycosphere, input CAS, and negative controls (**Fig. 2b**). However, the separation was smaller for fungi than for bacteria (18.1% of variance in the fungal composition, *P* <0.001, versus 26.1% in bacterial composition, *P* <0.001; **Fig. 2a, b**). Compared to the input CAS, Flavobacteriaceae and Lysobacteraceae were more abundant in the *C. reinhardtii* phycosphere (**Fig. 2c**, **Fig. S3**). Among fungi, three of the most abundant families found in input CAS (Pyronemataceae, Mortierellaceae and Trichosporonaceae) were depleted in the *C. reinhardtii* phycosphere (**Fig. 2d**, **Fig. S4**). Together, these results recapitulate previous observations for bacterial communities associated with *C. reinhardtii* and demonstrate that the model alga can also recruit specific soil fungal communities.

**Figure 2.**
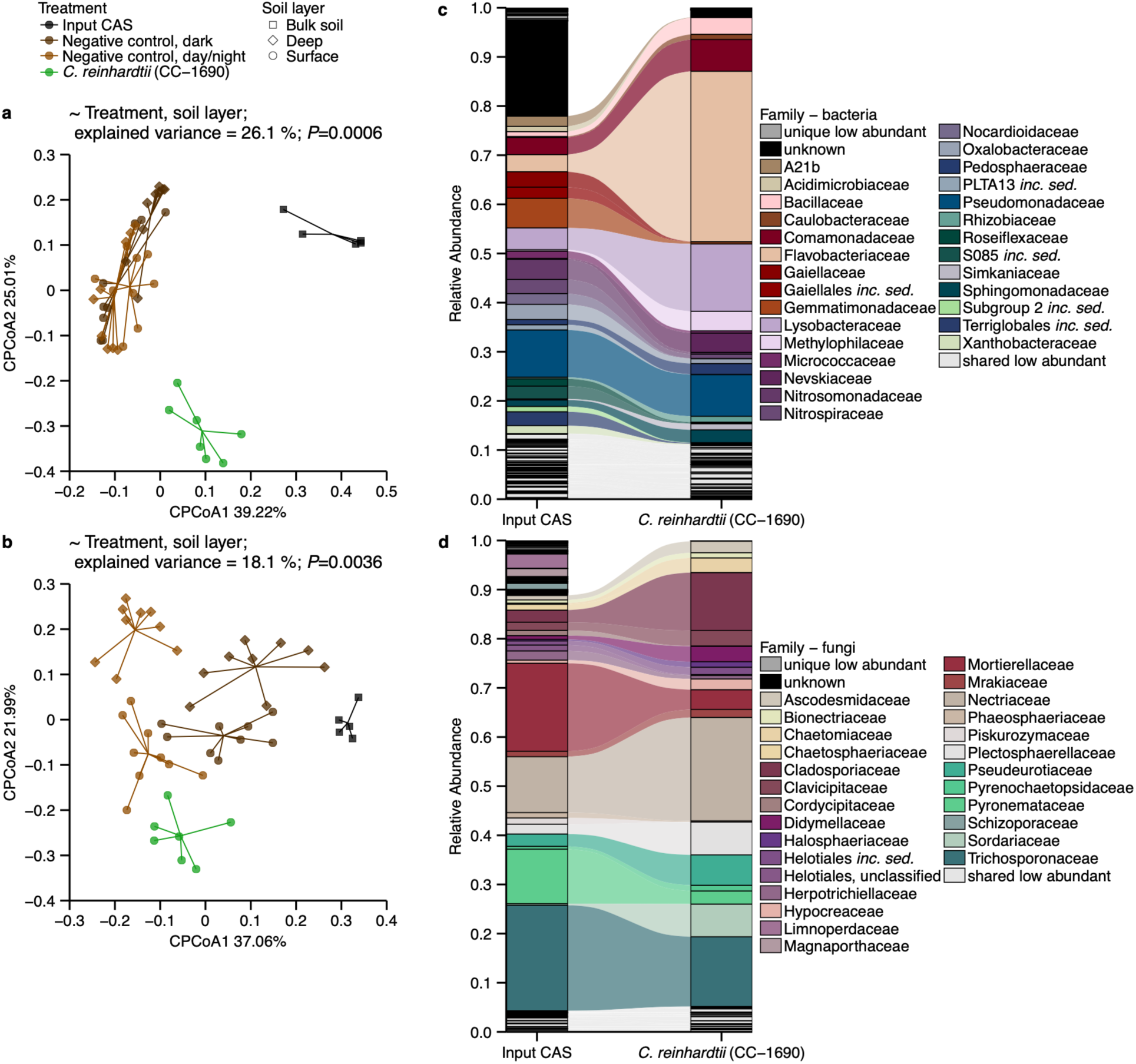
*C. reinhardtii* associates with bacteria and fungi derived from native CAS. (**a**, **b**) ASV-level beta-diversity analysis (Principal Coordinates Analysis of Bray-Curtis dissimilarities, constrained by treatment and soil layer combined as a single variable) of bacterial (a, 26.1% of the variance; *P* <0.001) and fungal communities (b, 18.1% of the variance; *P* <0.01) from the input CAS (*n* = 5), negative control, dark: deep soil layer (*n* = 6) and surface soil (*n* = 7), negative control, day/night: deep soil layer (*n* = 9) and surface soil (*n* = 9) and *C. reinhardtii* phycosphere (*n* = 5) samples. Points are color-coded based on the treatment, with point shapes corresponding to the soil layer. (**c**, **d**) Alluvial plots showing the average relative abundances of bacterial (**c**) and fungal (**d**) families in the input CAS samples and *C. reinhardtii* phycosphere samples.

### Subaerial algae reshape soil bacterial and fungal communities

To test whether the ability to assemble phycosphere microbiota from soil-borne bacteria and fungi is conserved across algal lineages, we inoculated CAS pots with individual strains from our taxonomically diverse subaerial algae collection, and characterized their bacterial and fungal phycosphere communities. Because *C. reinhardtii* associates with CAS-borne bacteria and fungi, this system provided a suitable framework to test whether other algae recruit soil microbes similarly to the model alga *C. reinhardtii*. During the experiment, substantial algal growth was visible on the soil surface, which was confirmed by extracting and measuring total chlorophyll content in surface soil samples (**Fig. S5a**). Samples from the algae-inoculated pots were taken once algal growth was comparable to that of *C. reinhardtii*, or at the latest on day 35 (**Materials and Methods**).

In line with our hypothesis, analysis of beta-diversity using CPCoA showed that bacterial communities were separated according to algal presence along the first axis, and by host taxonomy along the second (**Fig. 3a**, **Fig. S6a**). We observed that treatment (host identity and soil layer) explained the largest variance proportion (29.6%, *P* <0.001), while light regime and total chlorophyll content explained much smaller, non-significant fractions of variance (3.7% and 4.3%, n.s.).

**Figure 3.**
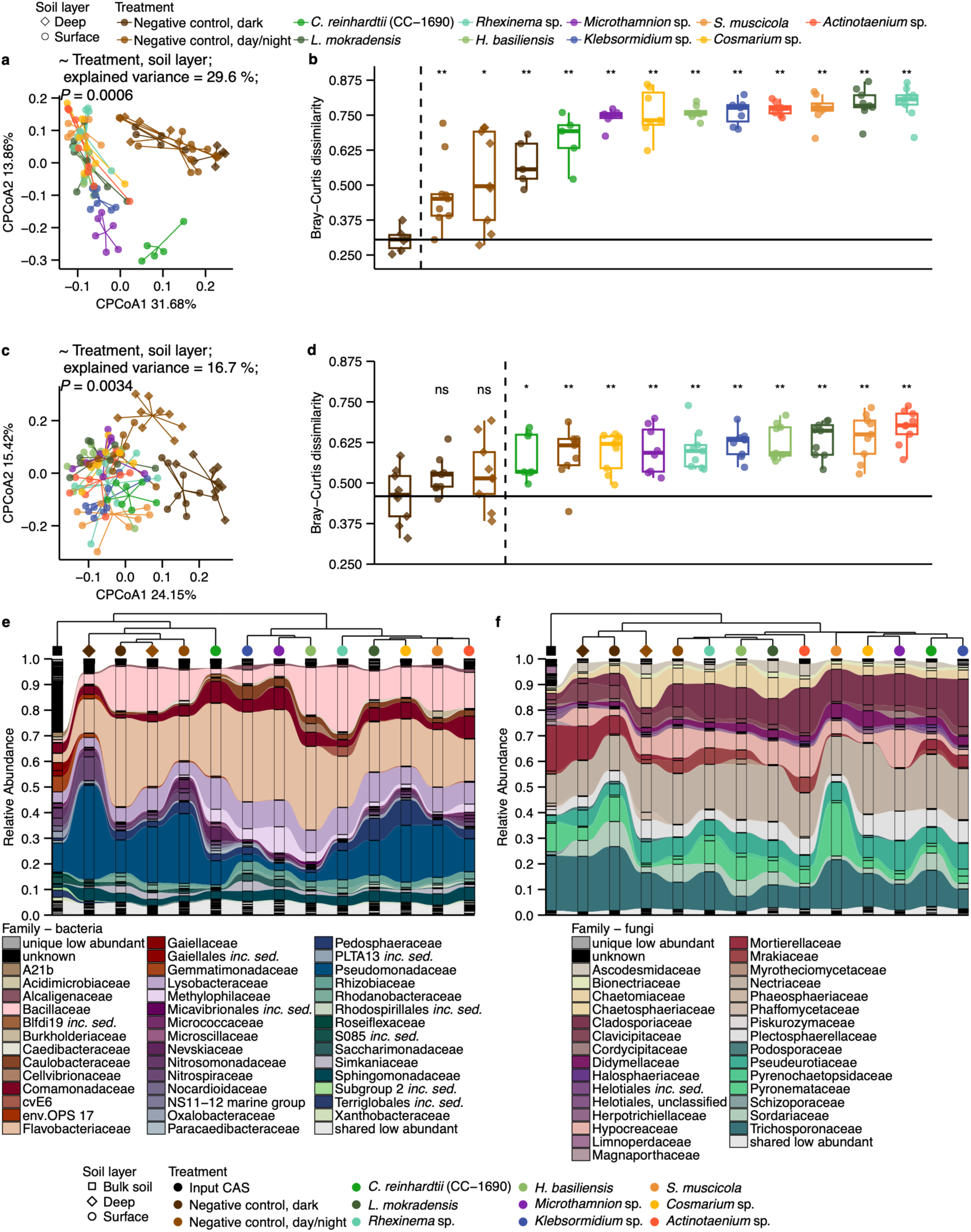
Host taxonomy influences bacterial community structure in soil, and to a lesser extent, the fungal community. **(a)** ASV-level beta-diversity analysis (Principal Coordinates Analysis of Bray-Curtis dissimilarities, constrained by Treatment and soil layer combined as a single variable, 29.6% of the variance; *P* <0.001) of bacterial communities from the negative control samples and phycosphere samples. (**b**) ASV-level analysis of Bray-Curtis dissimilarities of bacterial communities from negative control and phycosphere samples, compared to the averaged negative control, dark, deep soil sample. Significant differences are marked with stars (two-sided Mann-Whitney test, with Benjamini-Hochberg correction; ns - not significant, *P* >0.05, *− 0.05 ≥*P *>0.01, − 0.01 ≥*P *>0.001,* - 0.001 ≥*P* >0.0001, - *P* <0.0001), and samples are color-coded based on the treatment, with point shapes corresponding to the soil layer. The dashed vertical line separates the experimental conditions not significantly different from the averaged negative control, dark, deep soil from those significantly different, and the horizontal line shows the average distance of the individual negative control, dark, deep soil samples to the averaged one. (**c**) ASV-level beta-diversity analysis (Principal Coordinates Analysis of Bray-Curtis dissimilarities, constrained by Treatment and soil layer combined as a single variable, 16.7% of the variance; *P* <0.01) of fungal communities from the negative control samples and phycosphere samples. (**d**) ASV-level analysis of Bray-Curtis dissimilarities of fungal communities from negative control and phycosphere samples compared to the averaged negative control, dark, deep soil sample. Significant differences were analyzed and are marked like in panel (**b**). (**e**, **f**) Alluvial plots showing the overlap between bacterial (**e**) and fungal (**f**) communities of native CAS, negative controls and algal phycospheres. The dendrograms show the result of ASV-level analysis of Bray-Curtis dissimilarities of average community composition (per Treatment and soil layer) compared by hierarchical clustering. *n* per Treatment and soil layer between 5 and 9.

In order to assess differences in the capacity of the algae to recruit distinct bacterial communities from the surrounding soil, we quantified amplicon sequence variants (ASV) level Bray-Curtis dissimilarities between the phycosphere samples of each host relative to negative controls (deep soil compartment kept in the dark; **Fig. 3b**). This analysis revealed strong divergence of all phycosphere samples from the controls, while the separation between the different controls was significant, but less pronounced. Similar significant differences were observed when comparing phycosphere samples to a reference soil surface control (day/night incubation), further supporting a host-dependent effect on phycosphere bacterial community assembly (**Fig. S5b**). In both comparisons, we did not observe grouping according to algal division (Chlorophyta and Streptophyta). Although we detected a host-dependent effect on the soil bacterial community, the ASV-level Bray-Curtis dissimilarities did not correlate significantly with algal phylogenetic distances (Mantel Spearman *r* = 0.10, *P* = 0.23; **Supplementary Table 2**).

Fungal communities showed a similar, but less pronounced response to the presence of the different algae (**Fig. 3c**). In contrast to bacterial communities, clustering along the first axis in beta diversity was primarily driven by light regime (**Fig. 3c**). However, fungal samples also separated according to host taxonomy along the second axis (**Fig. 3c**), indicating that the algal host also shapes fungal community structure. This host-dependent effect was of similar magnitude for Streptophyta and Chlorophyta (17.4% and 16% of variance) (**Fig. S6b**), and within Chlorophyta, the model *C. reinhardtii* phycosphere samples were further differentiated from those associated with other Chlorophyta derived from the Eifel natural site. This was also true for bacterial communities, suggesting that host origin might influence the phycosphere community composition (**Fig. S6**).

Similarly to bacterial communities, we computed fungal ASV-level Bray-Curtis dissimilarities to the reference negative control, which revealed significant differences for all phycosphere samples but not for the other negative controls (**Fig. 3d**). However, unlike for bacterial communities, algal presence did not universally increase fungal community dissimilarity compared to the negative control samples from surface soil exposed to light (**Fig. S5c**), a separation that was only detected for two Streptophyta hosts, *Klebsormidium* sp. and *S. muscicola* (**Fig. S5c**). Consistently with these observations, the ASV-level Bray-Curtis dissimilarities did not correlate significantly with algal phylogenetic distances (Mantel Spearman *r* = 0.13, *P* = 0.20, **Supplementary Table 2**).

To identify taxa driving compositional changes, we examined family-level RA shifts between phycosphere and negative control communities (**Fig. 3e-f**). Among bacteria, Caulobacteraceae, Rhizobiaceae and Methylophilaceae were consistently more abundant in the phycospheres than in the controls, while Pseudomonadaceae showed host-dependent variation, with some ASVs moderately changing in RA and others being excluded from the phycospheres (ASV-level enrichment analysis, one-sided Mann-Whitney test, BH-adjusted *P* <0.05, **Materials and Methods**; **Fig. S7, S8, S9**). In contrast, fungal communities clustered primarily by light regime with no clear separation between phycosphere and uninoculated controls (**Fig. 3f**). The fungal families dominating in CAS, Mortierellaceae, Trichosporonaceae, Pyronemataceae, markedly declined in the phycospheres, while Cladosporiaceae increased 4.1-to 7.5-fold (**Fig. S10, S11**). Additionally, taxa belonging to the core microbiota of photosynthetic organisms, such as Hypocreaceae (Hypocreales) and Pyrenochaetopsidaceae (Pleosporales) were present in the phycospheres and increased in abundance compared to the initial fungal community in CAS (Durán, 2024, Garrigues *et al.,* 2026).

Together, these results show that patterns observed for *C. reinhardtii* are found in other subaerial algae and that algal hosts strongly modified soil bacterial communities while exerting a comparatively weaker effect on fungal communities.

### Host taxonomy is associated with phycosphere composition in the liquid system

Given our earlier observations in the soil-based system, we then asked whether the host effect persists, and if a phylogenetic signal emerges when environmental heterogeneity is reduced. To this end, we performed experiments in the mesocosm system with carbon-free medium (Durán *et al*., 2022), which allows controlled observation of microbial community assembly over time. Since the growth medium contains no external organic carbon source, the algal exudates are the sole organic carbon source for the bacterial community.

As with the soil-based system, we first analyzed the phycosphere community of *C. reinhardtii.* Along 35 days *C. reinhardtii* established a distinct phycosphere community, clearly separating from the input CAS-derived bacteria and the negative controls (**Fig. 4a**). The taxonomic composition of the community progressively changed over time (**Fig. 4b**). Pseudomonadaceae became increasingly depleted in the phycosphere relative to controls, while Xanthobacteraceae nearly disappeared by day 14. Conversely, Sphingomonadaceae moderately increased, and Flavobacteriaceae and Weeksellaceae were detected exclusively in the phycosphere samples at the end of the experiment (**Fig. 4b**, **Fig. S12**). Together, these results confirmed that in the liquid system, *C. reinhardtii* assembles a distinct phycosphere community that undergoes compositional changes over time. This progressive differentiation of the bacterial communities, combining selective elimination of some taxa and host-specific enrichment of others, suggests active filtering by *C. reinhardtii* rather than stochastic drift.

**Figure 4.**
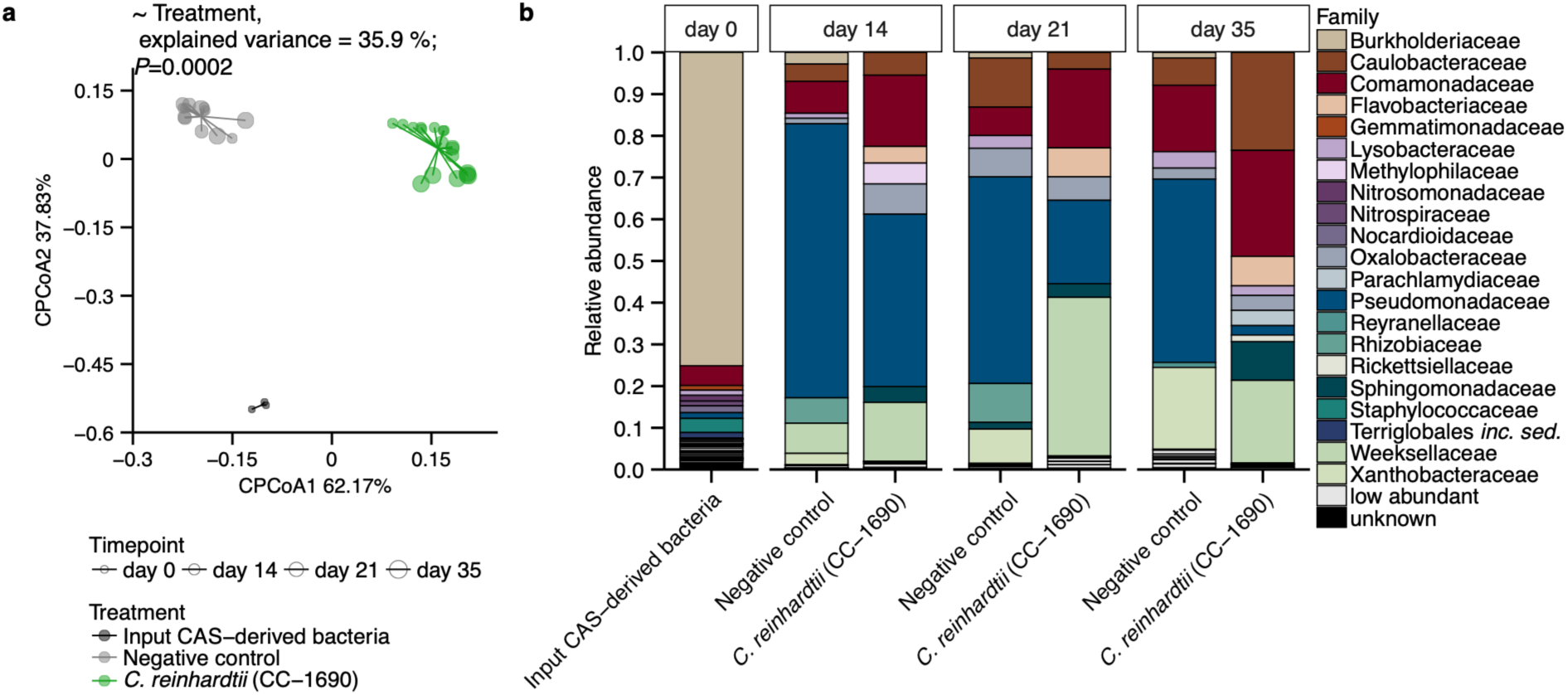
*C. reinhardtii* establishes a unique phycosphere bacterial community from CAS-derived bacteria in a liquid-based system. (**a**) ASV-level beta-diversity analysis (Principal Coordinates Analysis of Bray-Curtis dissimilarities, constrained by treatment, 35.9% of the variance; P <0.001) of bacterial communities from samples obtained from a liquid-based system. The samples are color-coded based on the treatment: input CAS-derived bacteria (black, n = 3), negative control (gray; n = 5 for day 14 and 21 and n = 3 for day 35) and *C. reinhardtii* (green; n = 8 for day 14, n = 4 for day 21 and n = 6 for day 35) and the point size corresponds to the timepoint. (**b**) Barplots of the average relative abundances of bacterial families in the input CAS-derived bacteria samples, negative control samples and C. reinhardtii phycosphere samples.

Consistent with observations for *C. reinhardtii* and the soil system experiment, all algae assembled distinct phycosphere communities (**Fig. 5a**). Interestingly, in the comparison of phycosphere amplicon profiles, samples from Chlorophyta and Streptophyta algae generally separated along the first ordination axis and clustered by host taxonomy along the second, indicating a strong influence of the host on the bacterial community composition (**Fig. 5a**). In contrast, we observed little separation by timepoint, suggesting that for several hosts phycosphere community composition had largely stabilized by day 14 (**Fig. 5a**). At the ASV level, phycosphere communities differed strongly and significantly from the no-alga control, with mean Bray-Curtis dissimilarities ranging from 60% to 95% across algal host (**Fig. 5b**). Moreover, ASV-level Bray-Curtis dissimilarities were correlated with algal phylogenetic distances (Mantel test, Spearman’s *r* = 0.66, *P* = 0.008; **Supplementary Table 2**), supporting the importance of the host identity for the bacterial community structure and indicating a phylosymbiotic signal under spatially unstructured conditions.

**Figure 5.**
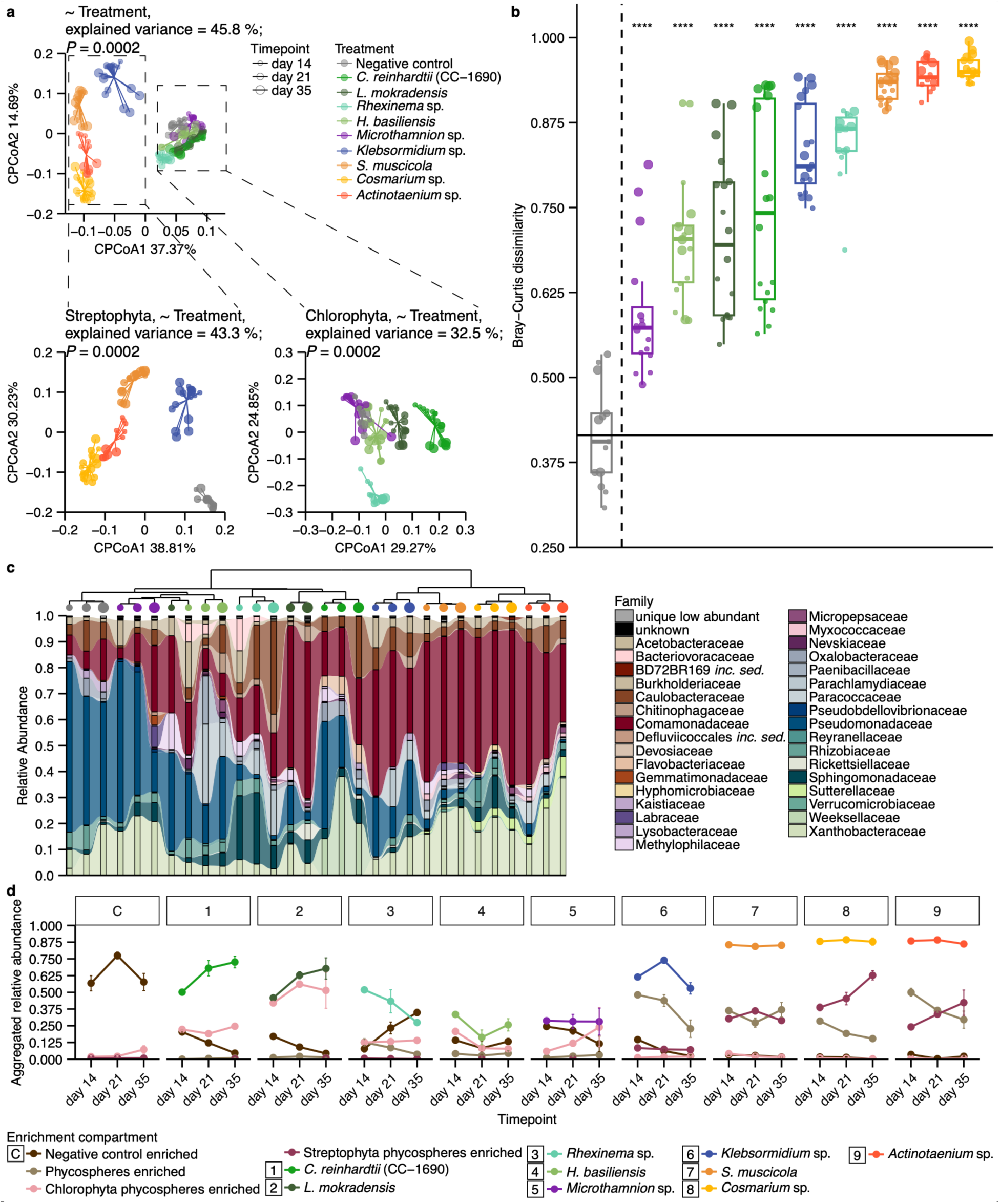
Phycosphere bacterial community depends on the algal host taxonomy. (**a**) ASV-level beta-diversity analysis of bacterial communities from samples obtained from a liquid-based system, (per treatment and timepoint, *n* between 3 and 9, Principal Coordinates Analysis of Bray-Curtis dissimilarities constrained by Treatment, 45.8% of the variance; *P* <0.001). Samples are color-coded by Treatment, with point sizes corresponding to the timepoint. The dashed lines show separate analyses for negative control and Streptophyta (left, 43.3% of the variance; *P* <0.001) or Chlorophyta (right, 32.5% of the variance; *P* <0.001). (**b**) ASV-level analysis of Bray-Curtis dissimilarities of bacterial communities from negative control and phycosphere samples, compared to the averaged negative control sample. Significant differences are marked with stars (two-sided Mann-Whitney test, with Benjamini-Hochberg correction; ns -not significant, *P* >0.05, *- 0.05 ≥*P *>0.01, - 0.01 ≥*P *>0.001,* - 0.001 ≥*P* >0.0001, - *P* <0.0001), and samples are color-coded based on the Treatment, with point sizes corresponding to the timepoint. The dashed vertical line separates the Treatments not significantly different from the averaged negative control from those significantly different, and the horizontal line shows the average distance of the individual negative control samples to the averaged one. (**c**) Alluvial plot showing the overlap between bacterial communities of negative controls and algal phycospheres. The dendrograms show the result of ASV-level analysis of Bray-Curtis dissimilarities of average community composition (average per treatment and timepoint) grouped by hierarchical clustering. (**d**) Dynamics of relative abundances of ASVs enriched over time in negative controls, all phycospheres, Streptophyta or Chlorophyta phycospheres and single alga phycospheres, color-coded by the enriched compartment. The plot is faceted by treatment: C - negative control, 1 - *C. reinhardtii* (CC-1690), 2 - *L. mokradensis*, 3 - *Rhexinema* sp., 4 - *H. basiliensis*, 5 – *Microthamnion* sp., 6 - *Klebsormidium* sp., 7 - *S. muscicola*, 8 - *Cosmarium* sp., 9 - *Actinotaenium* sp.

Temporal dynamics of phycosphere assembly, assessed based on the changes in Bray-Curtis dissimilarities, varied between hosts. Zygnematophyceae established distinct communities early on, whereas phycosphere communities of the Chlorophyceae (*C. reinhardtii*, *L. mokradensis*) continued to change over time (**Fig. 5b**). For some hosts (*S. muscicola* and the two Desmidiales algae, *Actinotaenium* sp. and *Cosmarium* sp., all Zygnematophyceae), the Bray-Curtis dissimilarities, which show the extent to which the phycosphere community differs from the control, were already high on day 14, indicating these algae select for specific microbes faster. While *S. muscicola*, *Actinotaenium* sp. and *Cosmarium* sp. showed high Bray-Curtis dissimilarities from the negative controls already from day 14 onwards, by day 35 three other algae (*L. mokradensis*, *C. reinhardtii*, *Klebsormidium* sp.) reached comparable values, while for the other three (*Rhexinema* sp., *H. basiliensis* and *Microthamnion* sp.), the dissimilarities were lower. These observations indicate that the extent to which algae modify the bacterial community, as well as the temporal dynamics of the phycosphere community assembly, differ between hosts and are not more similar within than between host taxonomic divisions, even though community composition itself separates by division (**Fig. 5a**).

We then sought to identify the bacterial taxa driving the compositional changes in the communities. We first analyzed RA changes and observed that Pseudomonadaceae dominated negative control samples, but were depleted in the phycospheres of *S. muscicola*, *Klebsormidium* sp., *Cosmarium* sp. and *Actinotaenium* sp., where Comamonadaceae were up to 169-fold more abundant than Pseudomonadaceae (**Fig. 5c**, **Fig. S13**, **S14**). The depletion of Pseudomonadaceae in the phycospheres was consistent with the observations in the soil system, suggesting that despite differences between the experimental systems, some patterns in the phycosphere community assembly are generalizable (**Fig. 3**, **Fig. 5**, **Fig. S15**).

To further characterize the bacterial taxa driving the differences between the negative control and the phycospheres, we performed an enrichment analysis (one-sided Mann-Whitney test, BH-adjusted *P*<0.05). Enrichment analysis identified ASVs enriched in the negative control, all phycospheres, phycospheres of Chlorophyta or Streptophyta hosts, and individual host-specific ASVs (**Fig. 5d**, **Fig. S16**). ASVs enriched in the negative control accounted for 57-78% RA in the negative control samples, but below 35% in the phycospheres (**Fig. 5d**). Only four ASVs were enriched in all algal hosts, suggesting only a minimal shared core of bacteria associating with all the investigated algae at this taxonomic level. Some ASVs were consistently enriched per algal division, indicating algal-division similarities rather than a large universal phycosphere microbiota. Generally, Chlorophyta-enriched ASVs represented 6-56% RA in Chlorophyta phycospheres but less than 5% in Streptophyta phycospheres. Conversely, Streptophyta-enriched ASVs accounted for 7-63% RA in Streptophyta phycospheres but were almost absent from the Chlorophyta phycospheres. Mirroring the variation in temporal dynamics of Bray-Curtis dissimilarities (**Fig. 5b**), the temporal trajectories of the host-enriched ASVs also differed among hosts. Some algae (*S. muscicola*, *Cosmarium* sp., and *Actinotaenium* sp.) established stable phycosphere communities early on, suggesting strong host influence on the microbial community. In contrast, Chlorophyceae (*C. reinhardtii*, *L. mokradensis*) showed a gradual increase in host-enriched ASVs over time, reaching up to 73% RA. Other algal hosts displayed a more variable or declining pattern. The observed patterns in community composition, together with the ASV-level Bray-Curtis dissimilarities correlating with algal phylogenetic distances (Mantel Spearman *r* = 0.66, *P* = 0.008; **Supplementary Table 2**), indicate a strong phylogenetic signal in community composition. However, as algal hosts differed in growth rates and biomass accumulation, we cannot fully disentangle phylogenetic effects from differences in resource availability.

Together, these results show that, in the liquid system, where bacterial growth is entirely driven by algal-derived organic carbon, host phylogeny is correlated with the phycosphere composition, and community dissimilarity mirrored host phylogeny. The host effects detected in soil were therefore not an artefact of that system but were amplified once environmental spatial heterogeneity was removed, with differences in the speed at which each host assembled its community.

## Discussion

Microscopic algae are abundant in soils (Jassey *et al*., 2022), yet most of what is known about the phycosphere microbiota comes from aquatic systems and, on land, from the bacterial communities associated with *C. reinhardtii* and a few other species (Seymour *et al*., 2017; Durán *et al*., 2022). Here, we show that nine subaerial green algae spanning five classes of Chlorophyta and Streptophyta each assemble a distinct phycosphere community from the same pool of soil microbes, in natural soil as well as in a liquid system in which algal photosynthates are the only source of organic carbon. This host effect extended to soil-borne fungi, although it was weaker than for bacteria.

The presence of this capacity to assemble a multi-kingdom microbiota in both divisions of the green lineage raises the possibility that it was already present in their common ancestor and was retained during the colonization of land. In that case, the preadaptation for symbiosis proposed for the algal ancestor of land plants (Delaux *et al*., 2015) would not be limited to the symbiosis signaling pathway and would include a more general ability to structure a microbial community. Alternatively, host-specific communities could arise without any trait dedicated to the recruitment of microbes, for instance, as a consequence of each alga releasing a different mixture of organic compounds. In the absence of a living host, bacterial communities assembled on mixtures of phytoplankton metabolites can be predicted from those assembled on the individual compounds (Fu *et al*., 2020), and communities from different inocula converge at the family level when grown on the same carbon source (Goldford *et al*., 2018). Our observations are compatible with both scenarios. Only four ASVs were enriched in the phycosphere of all hosts in the liquid system, whereas most were specific to one division or to one host. Caulobacterales and Rhizobiales, which belong to the core microbiota of photosynthetic hosts (Durán *et al*., 2022; Garrigues *et al*., 2026), were more abundant in the phycosphere than in soil without algae, which could reflect a conserved host trait or the ability of these bacteria to grow on photosynthates from any host. In *C. reinhardtii*, however, bacterial communities separated from the alga by a membrane assemble differently from those in direct contact with it (Durán *et al*., 2022), indicating that diffusible carbon is not sufficient to explain phycosphere assembly. Our data therefore show that this capacity is shared and host-specific, but do not allow us to distinguish common ancestry from convergence.

In soil, host identity shaped bacterial and fungal communities, but community dissimilarity was not correlated with host phylogenetic distance. If the host traits that determine phycosphere composition are conserved along the host phylogeny, however, closely related algae would be expected to assemble more similar communities. This pattern, known as phylosymbiosis, does not require coevolution and can arise when host traits that filter the microbiota, such as exudate composition, covary with host relatedness (Mazel *et al*., 2018; Stock *et al*., 2022). We hypothesized that it is masked in soil, where organic matter provides carbon that is independent of the host and the soil matrix confines bacteria to small and poorly connected patches (Bäcker *et al*., 2026), so that only a fraction of the sampled community is exposed to the alga. If this is the case, a correlation with host phylogeny should emerge in the liquid system, where photosynthates are the only carbon source and the environment is not spatially structured. In line with this hypothesis, bacterial community dissimilarity was correlated with host phylogenetic distance in the liquid system, extending an earlier qualitative observation in five algae (Durán *et al*., 2022). A similar contrast exists in land plants, where the microbiota follows host phylogeny in roots and only weakly or not at all in the rhizosphere (Naylor *et al*., 2017; Fitzpatrick *et al*., 2018). Taken together, these observations suggest that the extent to which the microbiota reflects host phylogeny depends on how much of the available carbon is provided by the host.

The phycosphere microbiota of subaerial algae included fungi in addition to bacteria, similar to the root microbiota of land plants (Thiergart *et al*., 2020). However, bacterial communities were shaped by the identity of the host, whereas fungal communities were structured mainly by light regime. In land plants, fungal communities are likewise determined mainly by soil and site, and less by the host or the root compartment than bacterial communities (Bonito *et al*., 2014; Thiergart *et al*., 2020; Tkacz *et al*., 2020). The presence of the same asymmetry in algae, which lack roots and differentiated tissues, suggests that it originates from differences in the biology of bacteria and fungi, independently of the anatomy of the host. Bacteria take up soluble compounds of low molecular weight more efficiently than fungi (Wang and Kuzyakov, 2024), whereas a fungal mycelium extends over distances far larger than the phycosphere, so that the same individual can draw on resources outside it and is likely to be sampled in both the phycosphere and the surrounding soil. Notably, fungal communities differed significantly from those of illuminated soil without algae for only two hosts, both streptophytes, *Klebsormidium* sp. and *S. muscicola*, the latter belonging to the lineage most closely related to land plants. Streptophyte algae possess most components of the signaling pathway required for mycorrhizal symbiosis in land plants (Delaux *et al*., 2015), raising the possibility that this lineage has a greater capacity to interact with fungi. This effect was modest, and absolute quantification of fungal loads together with co-cultivation of streptophyte and chlorophyte algae with fungal isolates will be needed to test it.

Our conclusions are limited to nine hosts and to a single soil, which fixed the pool of microbes available for recruitment. Isolating bacteria and fungi from the phycosphere of each host would allow host-specific synthetic communities to be assembled and cross-inoculated, as previously done for *C. reinhardtii* and *A. thaliana* (Durán *et al*., 2022), to test whether phycosphere commensals preferentially colonize the host from which they were isolated (Wippel *et al*., 2021). Experiments in which the bacterial community is separated from each host by a membrane that allows the diffusion of metabolites would determine whether the requirement for physical proximity observed in *C. reinhardtii* is a general feature of phycosphere assembly.

Overall, our findings show that the assembly of a host-specific microbiota from soil, best known from the roots of land plants, also takes place around microscopic algae from both divisions of the green lineage. Given the abundance of microalgae in soils (Jassey *et al*., 2022), host-specific phycosphere communities may be a widespread and so far unrecognized source of microbial diversity at the soil surface.

## Materials and methods

### Algal phylogeny

Subaerial algae were initially identified based on their morphology, and their identity was confirmed by Sanger sequencing of 18S or rbcL sequences. DNA was extracted from the individual algal cultures either with the FastDNA™ SPIN Kit for Soil following the manufacturer’s instructions (MP Biomedicals, Solon, USA) or using the SPRI beads method. All fragments were amplified in reactions with 2 U SuperHotTaq DNA Polymerase, 1x incomplete buffer, 2 mM MgCl_2_ (Bioron GmbH, Ludwigshafen, Germany), 0.3% BSA, 0.2 mM dNTPs (ThermoFisher Scientific) and 0.3 μM primers. 18S fragment was amplified with primers EukA and EukB (Medlin *et al*., 1988) or CrN1F (Marin *et al*., 1998) and EukB. rbcL fragments were amplified with BP22-1F, BP23-724R (Fay *et al*., 1997), RH1 (Manhart, 1994) and 1385R (McCourt *et al*., 2000), or in a 2-step PCR with MaGo1F-MaGo-3R and MaGo2F-MaGo-3R (Gontcharov *et al*., 2004). All the primers used for amplification and sequencing are in Supplementary Table 3 and PCR programs are in Supplementary Table 4. The sequencing was performed by Eurofins Genomics, Germany. Low-quality bases were identified by visual inspection, and the sequences were trimmed in 4Peaks software. The trimmed high-quality sequences were assembled in UGENE (Okonechnikov *et al*., 2012) using function “Reads de novo assembly (with CAP3)” and blastn (2.14.1) was used to find best hits to confirm algal identity. Molecular identification results were compared with morphological observations, and, in some cases, an alternative taxonomic assignment was made based on the combined evidence (Supplementary Table 5).

### Algal culture conditions

*C. reinhardtii* was cultured in TP medium, pH 7.0 at 25 °C and 125 μmol m^−2^ s^−1^, either under continuous light with shaking on a rotatory shaker at 70 RPM, or, for the soil experiment, under a 16 h/8 h light/dark cycle without shaking. Subaerial microalgae (isolated as described in Durán *et al*., 2022) were cultured in modified Synthetic Freshwater Medium (SFM; Durán *et al*., 2022; Melkonian and Melkonian, 2026) under 12 h/12 h light/dark cycle without shaking. Cell growth was determined either by measuring samples in a Multisizer 4e Coulter counter (Beckman Coulter Inc., California, USA) particle counter with the Beckman Coulter Multisizer software (v4.03) or using an Infinite M200Pro (TECAN Austria GmbH, Grödig, Austria) plate reader with the TECAN i-control software (v2.0.10.0), to determine either absorbance at 750 nm or chlorophyll fluorescence (excitation 440/9 nm, emission 680/20 nm).

### Eifel site samples collection

Multiple samples were collected in June 2022 and July 2025 from soil close to the wet rock surface from which the subaerial algae were isolated (the Eifel site; Cheng *et al*., 2019; Durán *et al*., 2022). Multiple samples were taken from different spots. Samples were collected using a metal spatula wiped thoroughly with 70% ethanol, transported to the laboratory in a cooled box and stored at −20 °C until further processing.

### Soil experiment setup

6 cm × 6 cm plastic pots were filled completely with Cologne Agricultural Soil (CAS). Each CAS pot was placed on the lower part of a sterile Petri dish (90 mm diameter). Before inoculation, each Petri dish was filled with 25 mL sterile Milli-Q water to pre-moisturize the soil. *C. reinhardtii* cultures were centrifuged (1900 × *g*, 5 min, RT), washed three times with sterile 10 mM MgCl_2_ (1900 × *g*, 5 min, RT), and resuspended in 10 mL sterile 10 mM MgCl_2_. Cell density of the algal suspension was measured in the Multisizer 4e Coulter counter, diluted with sterile 10 mM MgCl_2_ to a cell density of 2.25 × 10^6^ cells/mL, and the chlorophyll fluorescence of the suspension was determined. Subaerial algae were centrifuged (800 × *g*, braking and acceleration speed 2, 15 min, RT), washed once with sterile 10 mM MgCl_2_ (same centrifugation settings), and resuspended in 30 mL sterile 10 mM MgCl_2_. Chlorophyll fluorescence of the algal resuspensions was determined, and if needed, the suspensions were diluted to a chlorophyll fluorescence of *C. reinhardtii* 2.25 × 10^6^ cells/mL suspension. Chlorophyll fluorescence as a proxy of algal culture cell densities was used as previously (Durán *et al*., 2022). If the fluorescence was lower, the suspensions were used directly to inoculate the pots.

Directly before the inoculation, soil in each pot was moistened with 10 mL sterile Milli-Q water, carefully applied with a pipette to ensure uniform moisture distribution. To inoculate, algal suspensions or sterile 10 mM MgCl_2_ (negative controls) were carefully applied to the soil surface using a pipette to ensure uniform distribution. The negative control, dark pots were covered with aluminum foil. The pots were placed on a tray, covered with a plastic cover with holes covered with parafilm, and transported to a climate chamber, where 25 mL sterile Milli-Q water were added to each Petri dish. The pots were incubated for up to 35 days under 16 h/8 h light/dark cycle. Soil moisture, algal growth, and general soil condition were assessed periodically. The position of the pots in the trays was shuffled weekly to minimize edge and location effects. Water was added to the Petri dishes under each pot or gently applied to the soil surface.

Samples from the algae-inoculated and control pots were taken either when the algae grew similarly to *C. reinhardtii* on day 21 (predominantly green soil surface) or at the latest on day 35, when some algal growth was observed in the control pots. Samples from the *C. reinhardtii* and *Klebsormidium* sp. were taken on day 21, *Microthamnion* sp. on day 28, and all the other algae and negative controls on day 35. Soil samples were collected using a sterile metal spatula. For each pot, a single spatula was used, and between taking replicate samples from each pot, it was washed with ethanol. In addition to soil samples for microbial profiling, surface soil samples were obtained for chlorophyll extraction as a proxy for the algal growth.

### Liquid-based experiment setup

Microbial soil wash was prepared as described previously (Durán *et al*., 2022). Briefly, 5 g CAS were manually resuspended in 30 mL of sterile 1x Tris-EDTA (TE) supplemented with 0.1% Triton X-100 (SERVA Electrophoresis GmbH, Heidelberg, Germany), homogenized by inversion in a rotary mixer (40 RPM, 30 min). Bigger soil particles were removed by centrifugation for 1 min at 450 × *g* and the supernatant was centrifuged again at 3000 × *g* for 20 min. Afterwards, the final microbial soil wash was prepared by resuspending the pellet in 50 mL of SFM. Subaerial algae pre-cultured for two weeks in SFM were inoculated to a similar chlorophyll fluorescence to that equivalent to *C. reinhardtii* CC-1690 culture with cell density of 10^5^ cells/mL in SFM. To each flask, 1 mL of the microbial soil wash was added, which equals ca. 10^9^ cells (estimate based on Multisizer 4e Coulter counter measurements). As a “negative control”, SFM inoculated with an estimate of 10^9^ cells from the microbial soil wash was used. The cultures were incubated for 35 days under 12/12 h light/dark cycle without shaking and samples for DNA extraction were collected at days 0, 7, 14, 21 and 35. Day 7 samples did not pass sequencing quality control and were excluded from all analyses.

### DNA extraction from soil and algal samples

Total DNA was extracted from the Eifel site and soil samples using the FastDNA™ SPIN Kit for Soil following the manufacturer’s instructions (MP Biomedicals, Solon, USA). DNA samples were eluted in 50 μL nuclease-free water and used for microbial community profiling after adjusting DNA concentration to 5 ng/µL.

### DNA extraction from liquid samples by alkaline lysis

Bacterial DNA from liquid samples was extracted using alkaline lysis, which we previously determined to be efficient for extracting DNA from liquid samples (Bai *et al*., 2015; Durán *et al*., 2022). Briefly, 12 μL of the sample were diluted in 20 μL of Buffer I (NaOH 25 mM, EDTA(Na) 0.2 mM, pH 12), mixed by pipetting, and incubated at 94 °C for 30 min. Next, 20 μL of Buffer II (Tris-HCl 40 mM, pH 7.46) was added to the mixture and the samples were stored at − 20 °C.

### SPRI beads preparation

To prepare SPRI beads for DNA extraction, SpeedBeads magnetic carboxylate modified beads (Cytiva 65152105050250) stock solution was added to a tube and placed on the magnet. After 5 min binding, the supernatant was discarded and the beads were washed twice with 1 volume 1x TE (10 mM Tris, 1 mM EDTA). The tube was removed from the magnet, the beads were mixed with 1 volume 1x TE, added to 50 volumes NaCl-PEG solution (Supplementary Table 6), and carefully mixed. After preparation, their performance was tested against commercial AMPure XP beads by mixing them with 0.5x GeneRuler 1 kb DNA Ladder (ThermoScientific^TM^) in different ratios, performing two washes with 80% ethanol, eluting, loading on 1% agarose gel and running for 20 min at 150 V. SPRI beads were stored in the dark at 4 °C until use.

### SPRI beads DNA extraction

Samples were ground in Bertin Precellys Evolution HP Homogenizator (7600 rpm, 20 s) three times, with snap freezing in liquid nitrogen between rounds. After grinding, the samples were shortly centrifuged at 4 °C and kept at −80 °C until grinding of all the samples was finished. To extract DNA, 800 µL of preheated lysis buffer (composition in Supplementary Table 6) (water bath, 55 °C, 20 min) was added to each sample, mixed thoroughly by hand and frozen in liquid nitrogen. Samples were ground again in Bertin Precellys Evolution HP Homogenizator (7600 rpm, 1 min) and incubated at 65 °C for 1 h. After incubation, they were mixed by inversion ∼20x and spun down (10000 × *g*, 5 min, RT). 500 µL of each sample was transferred to a deep well plate already filled with 187.5 µL 5 M potassium acetate. The plates were centrifuged at 3214 × *g* for 10 min, and 480 µL of each supernatant were transferred into clean plates. 288 µL SPRI beads (well mixed before, RT) were added to each well, mixed by inversion ∼20x, briefly spun down and incubated at RT for 5 min. The plate was placed on the magnet until the solution was clear. Supernatants were removed and each sample was washed twice with 80 % ethanol. The beads were air dried for 30 s, the plate was removed from the magnet and 20 µL nuclease-free water were added to each well. After mixing the solution well, the plate was placed back on the magnet and the clear solution was transferred to a new plate. The eluted DNA was used for microbial profiling.

### Chlorophyll extraction from soil samples

Soil samples were oven-dried (60 °C) in 15 mL-falcon tubes for several days until the soil weight in a given sample was constant, indicating the soil was fully dry. Chlorophyll extraction was performed by solvent extraction and shaking. Briefly, a spatula tip of CaCO3 (Carl Roth, Karlsruhe, Germany) was added to each soil sample, as well as 6 mL of DMSO (Sigma-Aldrich, Darmstadt, Germany). Samples were boiled for 90 min in a water bath (65 °C) and then shaken for 20 min on a horizontal shaker. After placing the tubes in an upright position and allowing the soil to settle, the supernatant was transferred to a new tube, and the procedure was repeated. Merged supernatants were centrifuged (10 min, 3000 × *g*, 15°C). Absorption (700 nm, 665 nm, 648 nm) was measured using the

Infinite M200Pro (TECAN Austria GmbH, Grödig, Austria) plate reader with the TECAN i-control software (v2.0.10.0), and total chlorophyll content was calculated according to the adapted formula Chl *a + b* [µg/g soil]=([(A665-A700) x 8.02 + (A648 -A700) x 20.2] x DF x S)/DSW, where DF -dilution factor, S -amount of solvent [ml], DSW -dry soil weight [g] (Caesar *et al*., 2018).

### 16S rRNA amplicon sequencing

16S rRNA amplicon sequencing from the liquid system experiment was performed as described previously (Durán *et al*., 2022). 16S rRNA from the Eifel site and soil samples was amplified using a two-step PCR amplification protocol. In the first step, V5-V7 region of the bacterial 16S rRNA was amplified in a 25 μL reaction volume containing 2 U SuperHotTaq DNA Polymerase, 1x incomplete buffer, 2 mM MgCl_2_ (Bioron GmbH, Ludwigshafen, Germany), 0.3% BSA, 0.2 mM dNTPs (ThermoFisher Scientific) and 0.3 μM primers 799F (Chelius and Triplett, 2001) and 1192R (Beckers *et al*., 2016) (sequences in Supplementary Table 3). PCR was performed (initial denaturation at 94 °C/2 min, followed by 25 cycles of 94 °C/30 s, 55 °C/30 s, 72 °C/30 s and final elongation at 72 °C/5 min). Afterwards, single-stranded DNA and proteins were digested by adding 1 µL Antarctic Phosphatase, 1 µL Exonuclease I and 3 µL Antarctic Phosphatase buffer (New England BioLabs GmbH, Frankfurt, Germany) to each sample. Reactions were incubated at 37 °C for 30 min, followed by enzyme deactivation (85 °C, 15 min) and centrifugation (3000 rpm, 10 min, 4 °C). 4.8 µL of the supernatant was used as the DNA template for a second PCR, prepared in the same way as described but reaction volume of 40 µL, primers including barcodes and Illumina adapters, and cycles reduced to 10.

PCR product concentration was assessed using PicoGreen^TM^ assay (Quant-iT^TM^ PicoGreen^TM^ dsDNA reagent, Life Technologies Corporation, ThermoFisher Scientific): 2 µL PCR product was mixed with 0.04 µL Pico 200x and 72.96 µL 0.5x TE, incubated for 2-5 min covered from light, and measured in Infinite M200Pro (TECAN Austria GmbH, Grödig, Austria) plate reader with the TECAN i-control software (v2.0.10.0) (excitation 485/9 nm, emission 535/20 nm). Samples were pooled based on their fluorescence intensities. The pooled sample was loaded on 1.5% agarose gel and run for 40 min, either at 130 V (medium-sized gel) or 90 V (small gel). Afterwards, the bands were excised and DNA was purified using the Gel and PCR-Clean-up kit (MACHEREY-NAGEL GmbH & Co. KG, Düren, Germany). DNA concentration was determined from fluorescence measurements with the QuantiFluor® dsDNA System (Promega GmbH) according to the manufacturer’s protocol. The quality assessment of the library and paired-end Illumina sequencing on NovaSeq PE250 platform was performed by Novogene GmbH.

### ITS2 spacer sequencing

ITS2 spacer from the Eifel site and soil samples was amplified according to a two-step PCR amplification protocol. In the first step, the ITS2 spacer region was amplified in a 10 µL reaction volume containing 2 U SuperHotTaq DNA Polymerase, 1x incomplete buffer, 2 mM MgCl_2_ (Bioron GmbH, Ludwigshafen, Germany), 0.3% BSA, 0.2 mM dNTPs (ThermoFisher Scientific) and 0.3 μM tagged primers ITS2_F1 and ITS2_R1 (Thiergart *et al*., 2020) (Supplementary Table 3). PCR was performed (initial denaturation at 94 °C/2 min, followed by 30 cycles of 94 °C/30 s, 59 °C/30 s, 72 °C/30 s and final elongation at 72 °C/5 min). Afterwards, single-stranded DNA and proteins were digested by adding 0.1 µL Antarctic Phosphatase, 0.05 µL Exonuclease I and 3.85 µL nuclease-free water to each sample. Reactions were incubated at 37 °C for 30 min, followed by enzyme deactivation (85 °C, 15 min) and centrifugation (3000 rpm, 10 min, 4 °C). 5 µL of the supernatant was used as the DNA template for a second PCR, prepared in the same way as described but reaction volume of 15 µL and primers including barcodes and Illumina adapters. PCR was performed (initial denaturation at 94 °C/2 min, followed by 10 cycles of 94 °C/30 s, 65 °C/30 s, 72 °C/30 s and final elongation at 72 °C/5 min).

PCR product concentration was assessed using PicoGreen^TM^ assay (Quant-iT^TM^ PicoGreen^TM^ dsDNA reagent, Life Technologies Corporation, ThermoFisher Scientific): 2 µL PCR product was mixed with 0.04 µL Pico 200x and 72.96 µL 0.5x TE, incubated for 2-5 min covered from light and measured in Infinite M200Pro (TECAN Austria GmbH, Grödig, Austria) plate reader with the TECAN i-control software (v2.0.10.0) (excitation 485/9 nm, emission 535/20 nm). Samples were pooled based on their fluorescence intensities. The pooled library was re-concentrated twice with AMPure XP beads (Beckman Coulter Inc.) and loaded on 1.5% agarose gel and run for 40 min, either at 130 V (medium-sized gel) or 90 V (small gel). Afterwards, the bands were excised and DNA was purified using the Gel and PCR-Clean-up kit (MACHEREY-NAGEL GmbH & Co. KG, Düren, Germany). DNA concentration was determined from fluorescence measurements with the Quantus^TM^ Fluorometer and QuantiFluor® dsDNA System (Promega GmbH) according to the manufacturer’s protocol. The quality assessment of the library and paired-end Illumina sequencing on NovaSeq PE250 platform was performed by Novogene GmbH.

### Analysis of 16S rRNA amplicon profiling

Amplicon sequencing data were processed in QIIME 2 2026.1 (Bolyen *et al*., 2019). Raw reads were imported, demultiplexed according to their barcode sequences, trimmed and paired reads were denoised with DADA2 within QIIME 2 to infer unique amplicon sequence variants (ASVs) after error correction and chimera removal (Callahan *et al*., 2016).

For taxonomic classification, a V5-V7 region-specific 16S rRNA gene classifier was trained using the SILVA 138.2 reference database (Quast *et al*., 2013; Yilmaz *et al*., 2014) and the RESCRIPt plugin for QIIME 2 (Bolyen *et al*., 2019). Briefly, full-length SSURef sequences were downloaded, RNA sequences were reverse-transcribed to DNA, and low-quality sequences containing ≥5 ambiguous bases or homopolymers of length ≥8 bp were excluded. Sequences were length-filtered by taxonomic domain with minimum lengths of 900 bp (Archaea), 1200 bp (Bacteria), and 1400 bp (Eukaryota) and dereplicated by identical sequence and taxonomy combinations (‘qiime rescript dereplicatè, uniq mode). The V5-V7 region was extracted with the primer pair AACMGGATTAGATACCCKG (forward) and ACGTCATCCCCACCTTCC (reverse), restricting fragment lengths to 250–450 bp. After additional dereplication to minimize redundancy, the naïve Bayes classifier was trained and evaluated on the dereplicated V5-V7 fragments (‘qiime rescript evaluate-fit-classifier’).

The trained classifier was used to assign taxonomy to the ASVs. Non-bacterial reads and ASVs detected exclusively in the initial axenic algal cultures were then removed, and the resulting abundance tables were imported into R (v4.4.0; R Core Team, 2024) for all downstream analyses.

### Analysis of ITS2 spacer amplicon profiling

The first round of amplicon sequencing data demultiplexing was performed by Novogene GmbH. Next, data were demultiplexed according to their barcode sequence using Cutadapt (Martin, 2011), and reads were merged and denoised with DADA2 within QIIME 2 (‘qiime dada2 denoise-singlè with ‘--p-trunc-qlen 0’ to retain full-length reads; Callahan *et al*., 2016). Unique amplicon sequencing variants (ASVs) were inferred from reads after error correction and chimera filtering. Taxonomic annotation of ASVs was performed using the naïve Bayesian classifier trained on UNITE data (Abarenkov *et al*., 2024). To train the classifier, reference sequences and taxonomy files were retrieved using the ‘qiime rescript get-unite-dat’ command with the parameters ‘--p-taxon-group eukaryotes’, ‘--p-cluster-id dynamic’, and ‘--p-singletons’ (Robeson *et al*., 2021). To improve classifier efficiency, SH accession identifiers were removed from the taxonomy strings. The full-length ITS naïve Bayes classifier was then trained and validated using ‘qiime rescript evaluate-fit-classifier’. To generate an abundance table, raw reads were mapped to the inferred ASVs. The resulting tables after excluding non-fungal reads were subsequently used for analyses of diversity and community compositions using R (v4.4.0; R Core Team, 2024).

### Statistics and reproducibility

For each experiment, the number of samples (*n*) is given in the figure legends. For comparisons, we used a two-sided, non-parametric Mann-Whitney test and when appropriate, *P*-values were adjusted for multiple comparisons using the Benjamini-Hochberg method. Whenever boxplots were used in figures, data were represented as median values (horizontal line), interquartile ranges (boxes, spanning Q1–Q3), and whiskers extending to the most extreme values within 1.5 × the interquartile range.

For the soil experiment, only samples with successful sequencing of both 16S (≥7200 bacterial reads) and ITS2 (≥1500 fungal reads) were kept for further analyses. In the soil experiment, a small subset of soil samples exhibited extreme dominance of the fungal order Pezizales, with relative abundances up to 86%. Although Pezizales were observed at moderate abundances in other samples across treatments, these extreme profiles formed a distinct subset and deviated from the overall community structure. Such dominance patterns might disproportionately influence multivariate analyses and obscure treatment-level effects. Because the pot constituted the experimental unit, all samples originating from three pots exhibiting such dominance were excluded collectively from downstream analyses. The exclusion criterion was based on the magnitude of Pezizales dominance (average Pezizales RA per pot 27.4%) rather than a fixed abundance threshold.

For analysis of alpha and beta diversities, rarefied ASV tables were used, while relative abundances were computed from non-rarefied data. Community structure was explored using principal coordinates analysis (PCoA) based on Bray-Curtis dissimilarities. For the soil experiment data, to quantify the association between environmental and experimental variables and microbial community composition, constrained analysis of principal coordinates (CAP; vegan::capscale) was applied. Each explanatory variable (Treatment × soil layer, algal class, algal presence/absence, light regime, and chlorophyll content) was evaluated in a separate single-factor model of the form: Bray-Curtis ∼ X. These models were interpreted descriptively to estimate associations between individual variables and community composition. Statistical significance tests on beta-diversity analyses were performed using anova.cca function from the *vegan* package (version 2.7-2; Oksanen *et al*., 2025) with 5000 permutations. The variance explained by each constraint was calculated as the ratio of the constrained inertia to the total inertia. To visualize treatment-associated community structure, CAP was performed with “Treatment, soil layer” as the constraining variable.

To isolate host-specific bacterial and fungal responses independent of environmental variation, analyses were restricted to phycosphere samples collected under identical conditions (surface soil, day/night light regime). Bray-Curtis dissimilarities were subset to the corresponding samples without recomputation. Host effects on fungal community composition were evaluated using PERMANOVA (vegan::adonis2) with algal treatment (individual alga/negative control) as the explanatory variable.

To assess whether microbial community similarity reflected algal evolutionary relatedness, we tested for phylosymbiosis using Mantel correlations between microbial Bray-Curtis dissimilarities and algal phylogenetic distances in R (v4.4.0; R Core Team, 2024) using package *ape* (5.8-1; Paradis and Schliep, 2019). Because marker-gene phylogenies (18S rRNA and rbcL) did not consistently resolve relationships among all algal strains and produced unstable topologies, we constructed a conservative, taxonomy-informed phylogenetic tree reflecting accepted higher-level relationships within

Chlorophyta and Streptophyta. Branch lengths were assigned using Grafen’s method (Grafen, 1989) and converted to an ultrametric tree to ensure comparability of phylogenetic distances. Phylogenetic distances were extracted as a cophenetic distance matrix and used exclusively for correlation analyses, without evolutionary inference. Microbial community dissimilarities were calculated using Bray-Curtis distances on group-averaged, rarefied ASV tables. Mantel tests were performed with 9,999 permutations using both Pearson and Spearman correlations to account for linear and monotonic relationships, respectively. Analyses were conducted separately for soil bacterial (16S), soil fungal (ITS), and liquid-culture bacterial (16S) communities. This approach has several limitations: the phylogenetic framework is based on taxonomy rather than sequence-derived branch lengths, and therefore reflects relative relatedness rather than evolutionary divergence times, and Mantel tests have limited statistical power at small sample sizes. Nevertheless, this analysis provides a conservative assessment of whether broad algal phylogenetic relatedness is associated with systematic differences in bacterial and fungal community composition across experimental contexts.

To identify bacterial ASVs significantly enriched in algal phycospheres relative to control treatments, or in control relative to phycospheres, we performed a series of one-sided Mann-Whitney tests on ASV relative abundances. ASVs were filtered to retain only those with a relative abundance ≥0.5 % in at least one sample, ensuring that low-abundance taxa did not inflate multiple-testing corrections. For each ASV, we tested whether its relative abundance was significantly more abundant than in the negative control samples in (i) each individual algal phycosphere compartment, (ii) all phycosphere samples combined, and (iii) subsets corresponding to Streptophyta or Chlorophyta hosts; and if it was significantly more abundant in the negative control than in all phycosphere samples combined (negative control enriched). *P*-values were adjusted using the Benjamini-Hochberg method and ASVs with adjusted *P*-values <0.05 were classified as significantly enriched.

### Transparency and AI use

Generative AI models (ChatGPT by OpenAI and Claude by Anthropic) were used to assist in drafting code templates, refining analytical workflows, language refinement and improving textual explanations. The models did not independently generate results or draw conclusions. All AI-generated content was reviewed, verified, and edited by the authors. All analyses were designed, executed, and interpreted by the authors. The authors retain full responsibility for the accuracy, originality, and integrity of the work.

## Acknowledgments

This work was supported by the European Research Council (ERC) under the European Union’s Horizon Europe research and innovation programme (PHYCOSPHERES, grant agreement No. 101077231) and the Cluster of Excellence on Plant Sciences (CEPLAS), funded by the Deutsche Forschungsgemeinschaft (DFG, German Research Foundation) under Germany’s Excellence Strategy (EXC 2048/1, project no. 390686111).

## Supplementary figures

**Figure S1.**
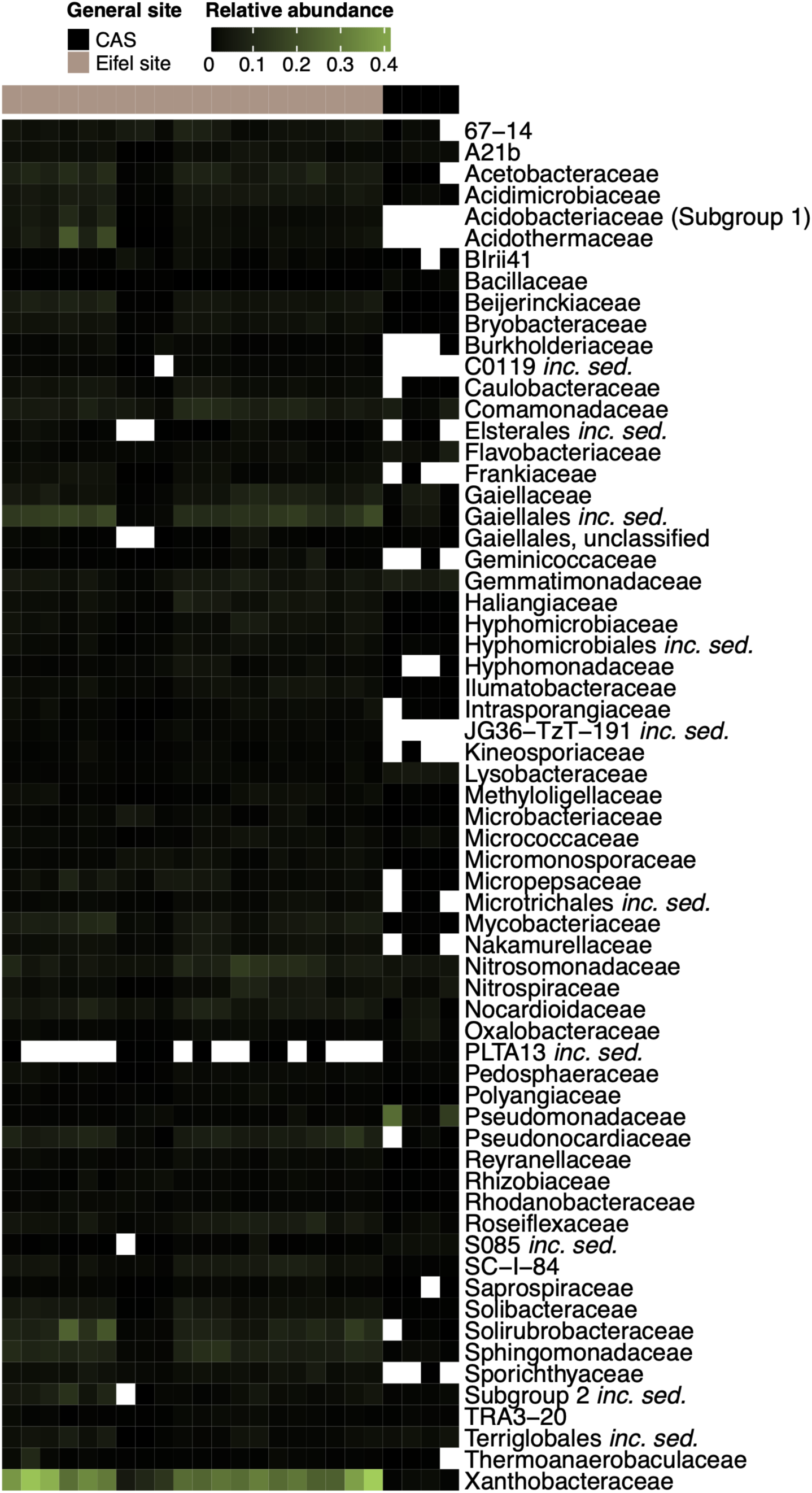
Heatmap of bacterial family-level relative abundances in samples from the Eifel site (*n* = 21) and CAS (*n* = 4). The heatmap shows relative abundances of bacterial families (rows) across samples (columns). Only families with ≥1% mean relative abundance in at least one site are included. Columns are annotated by site (top). Color intensity indicates relative abundance, with white representing zero abundance (family not detected in the sample).

**Figure S2.**
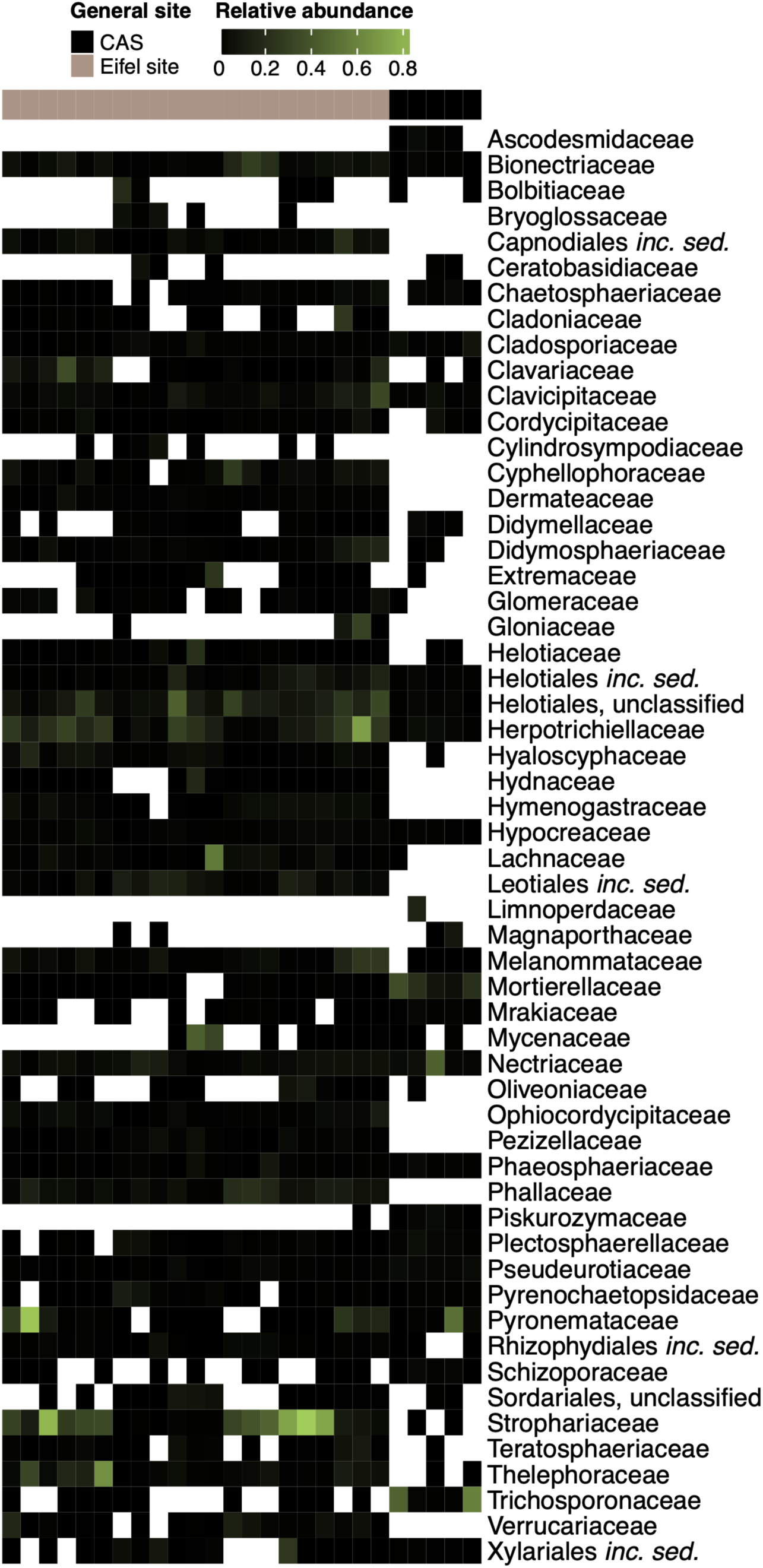
Heatmap of fungal family-level relative abundances in samples from the Eifel site (*n* = 21) and CAS (*n* = 5). The heatmap shows relative abundances of fungal families (rows) across samples (columns). Only families with ≥1% mean relative abundance in at least one site are included. Columns are annotated by site (top). Color intensity indicates relative abundance, with white representing zero abundance (family not detected in the sample).

**Figure S3.**
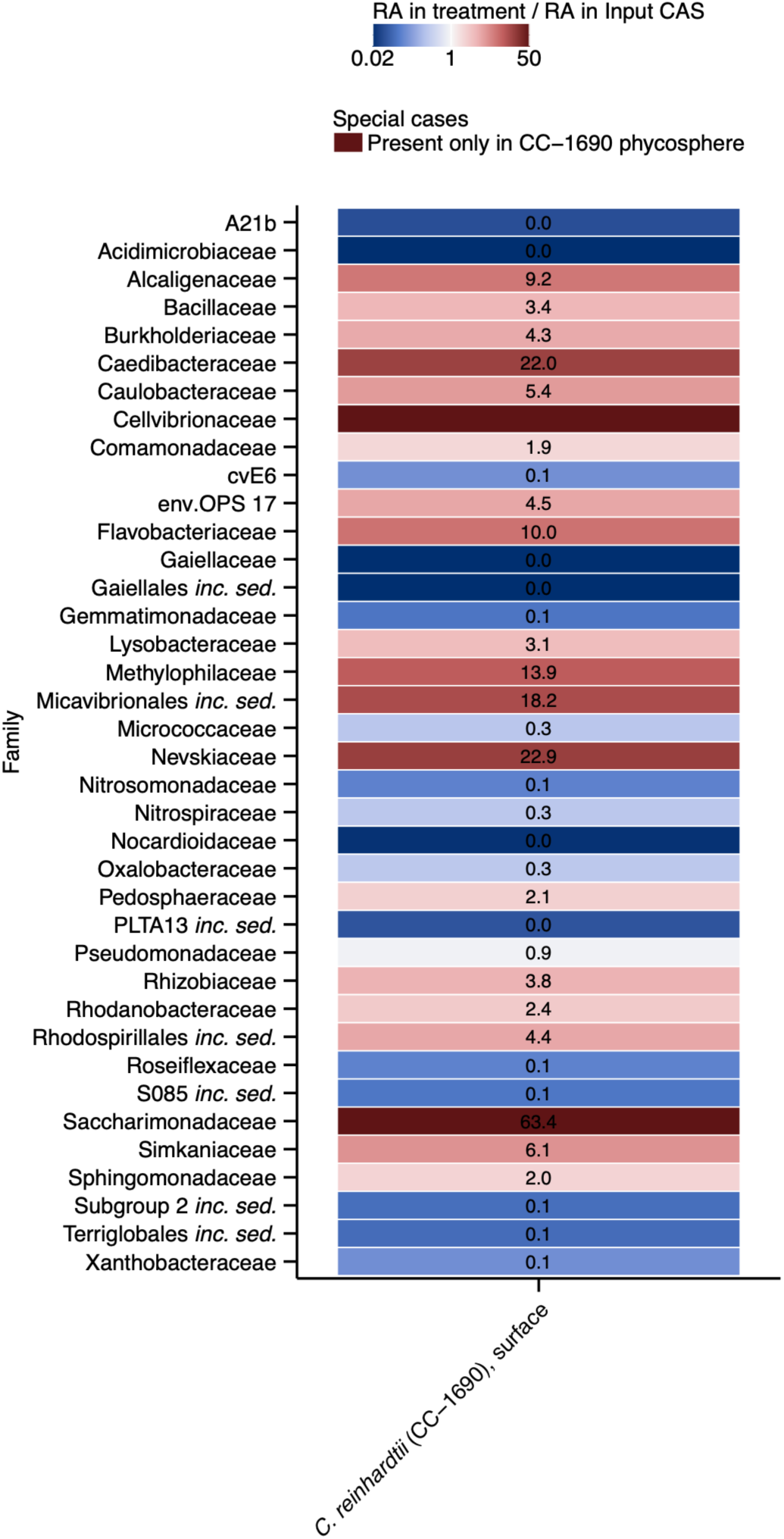
Ratios of mean family-level bacterial RA between *C. reinhardtii* CC-1690 phycosphere and input CAS (RA_treatment_/RA_input_). Cells are colored according to the ratio: shades of blue indicate a decrease in family-level RA in the phycosphere, and shades of red indicate an increase. Numbers represent the ratio value. Only families with ≥1% mean relative abundance in either input CAS or *C. reinhardtii* CC-1690 phycosphere are included.

**Figure S4.**
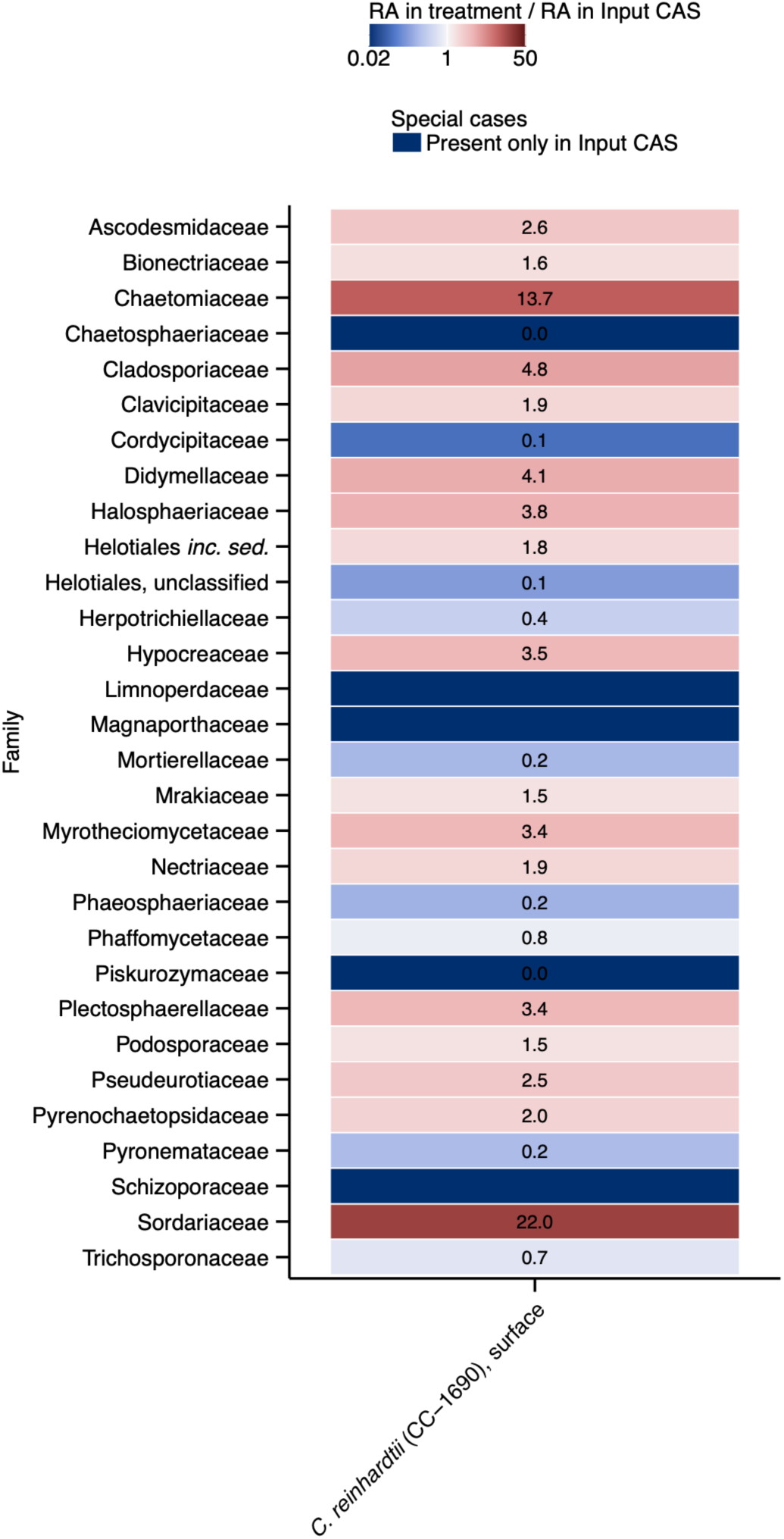
Ratios of mean family-level fungal RA between *C. reinhardtii* CC-1690 phycosphere and input CAS (RA_treatment_/RA_input_). Cells are colored according to the ratio: shades of blue indicate a decrease in family-level RA in the phycosphere, and shades of red indicate an increase. Numbers represent the ratio value. Only families with ≥1% mean relative abundance in either input CAS or *C. reinhardtii* CC-1690 phycosphere are included.

**Figure S5.**
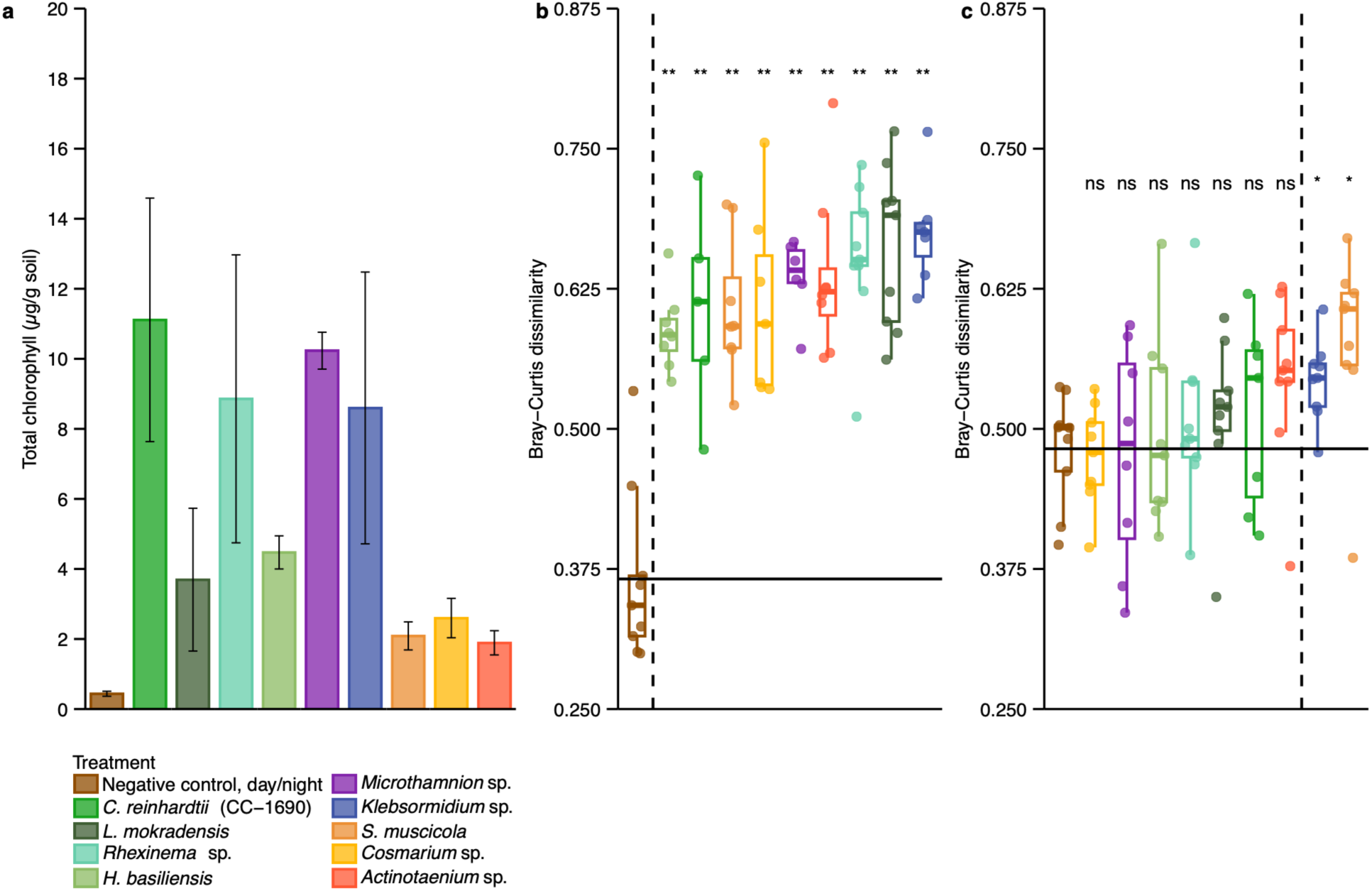
All algae grow on soil and significantly change soil bacterial community composition, while only some significantly influence the fungal community composition. (a) Total chlorophyll content in soil surface samples from pots inoculated with different algae and negative control. Bars represent mean total chlorophyll per gram of soil (± standard error, *n* = 3 biological replicates, each with 3 technical replicates) for each treatment. Colors correspond to individual algal treatments. The negative control incubated in the dark is excluded. (b) ASV-level analysis of Bray-Curtis dissimilarities of bacterial communities from negative control and phycosphere samples, compared to the averaged negative control, day/night, surface soil sample. Significant differences are marked with stars (two-sided Mann-Whitney test, with Benjamini-Hochberg correction; ns - not significant, *P* >0.05, *- 0.05 ≥*P *>0.01, - 0.01 ≥*P *>0.001,* - 0.001 ≥*P* >0.0001, - *P* <0.0001), and samples are color-coded based on the treatment, with point shapes corresponding to the soil layer. The dashed vertical line separates the experimental conditions not significantly different from the averaged negative control, day/night, surface soil from those significantly different, and the horizontal line shows the average distance of the individual negative control, day/night, surface soil samples to the averaged one. (c) ASV-level analysis of Bray-Curtis dissimilarities of fungal communities from negative control and phycosphere samples, compared to the averaged negative control, day/night, surface soil sample. Significant differences are marked as in panel (b) *n* per Treatment and soil layer between 5 and 9.

**Figure S6.**
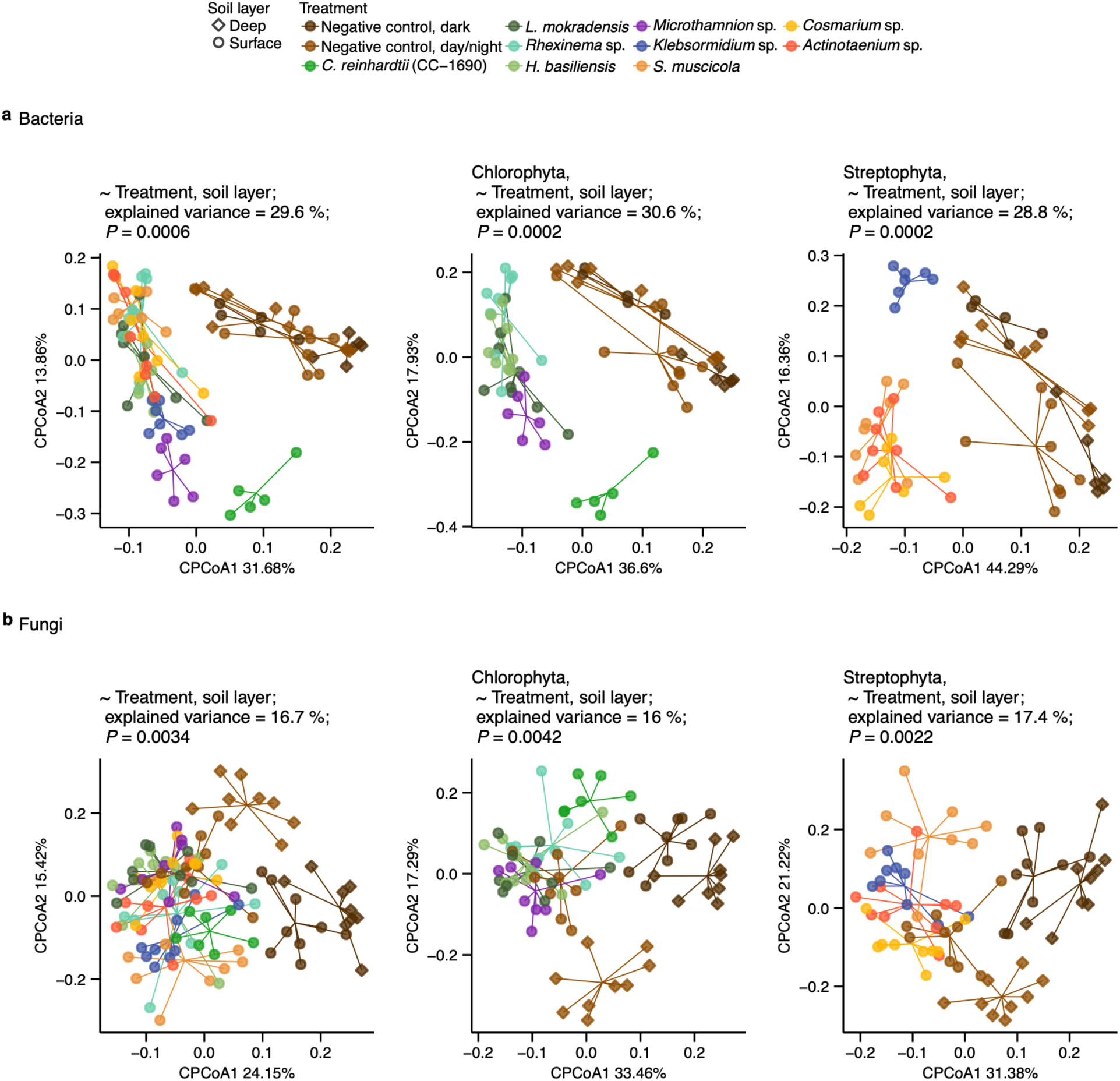
Phycosphere community structure depends on algal host taxonomy. (**a**) ASV-level beta-diversity analysis (Principal Coordinates Analysis of Bray-Curtis dissimilarities, constrained by treatment and soil layer combined as a single variable) of bacterial communities from the negative control samples and phycosphere samples, including an analysis of all treatments (29.6% of the variance; *P* <0.001; equivalent to Fig. 3) and separate analyses for negative control and Chlorophyta (30.6% of the variance; *P* <0.001) or Streptophyta (28.8% of the variance; *P* <0.001). (**b**) ASV-level beta-diversity analysis (Principal Coordinates Analysis of Bray-Curtis dissimilarities, constrained by Treatment and soil layer combined as a single variable) of fungal communities from the negative control samples and phycosphere samples, including an analysis of all treatments (16.7% of the variance; *P* <0.01) and separate analyses for negative control and Chlorophyta (16% of the variance; *P* <0.01) or Streptophyta (17.4% of the variance; *P* <0.01).

**Figure S7.**
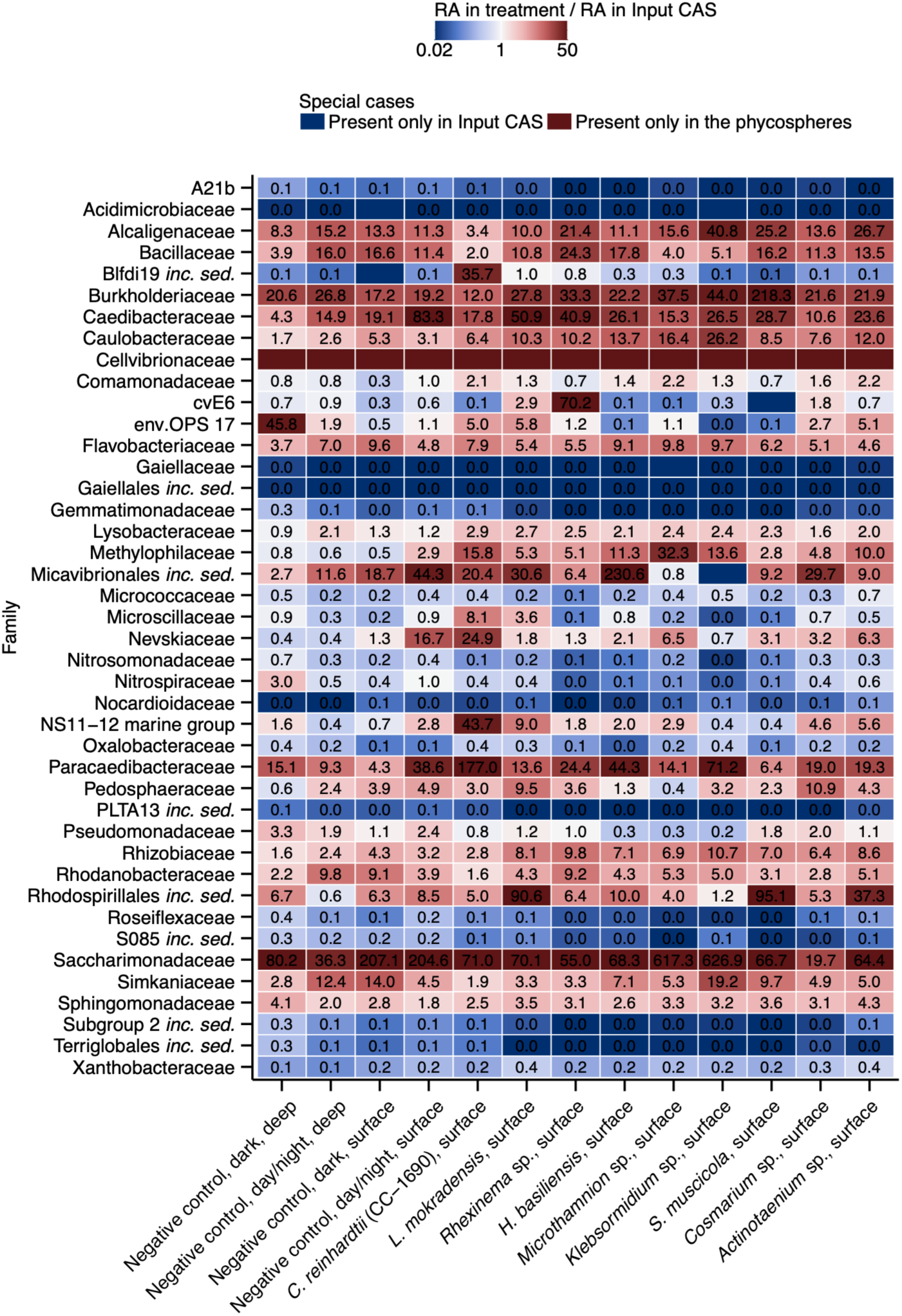
Ratios of mean family-level bacterial RA between phycospheres or negative controls and input CAS (RA_treatment_/RA_input_). Cells are colored according to the ratio: shades of blue indicate a decrease in family-level RA in the phycosphere, and shades of red indicate an increase. Numbers represent the ratio value; families detected only in the phycospheres or input CAS are shown without a number. Only families with ≥1% mean relative abundance in either input CAS or one of the treatments are included.

**Figure S8.**
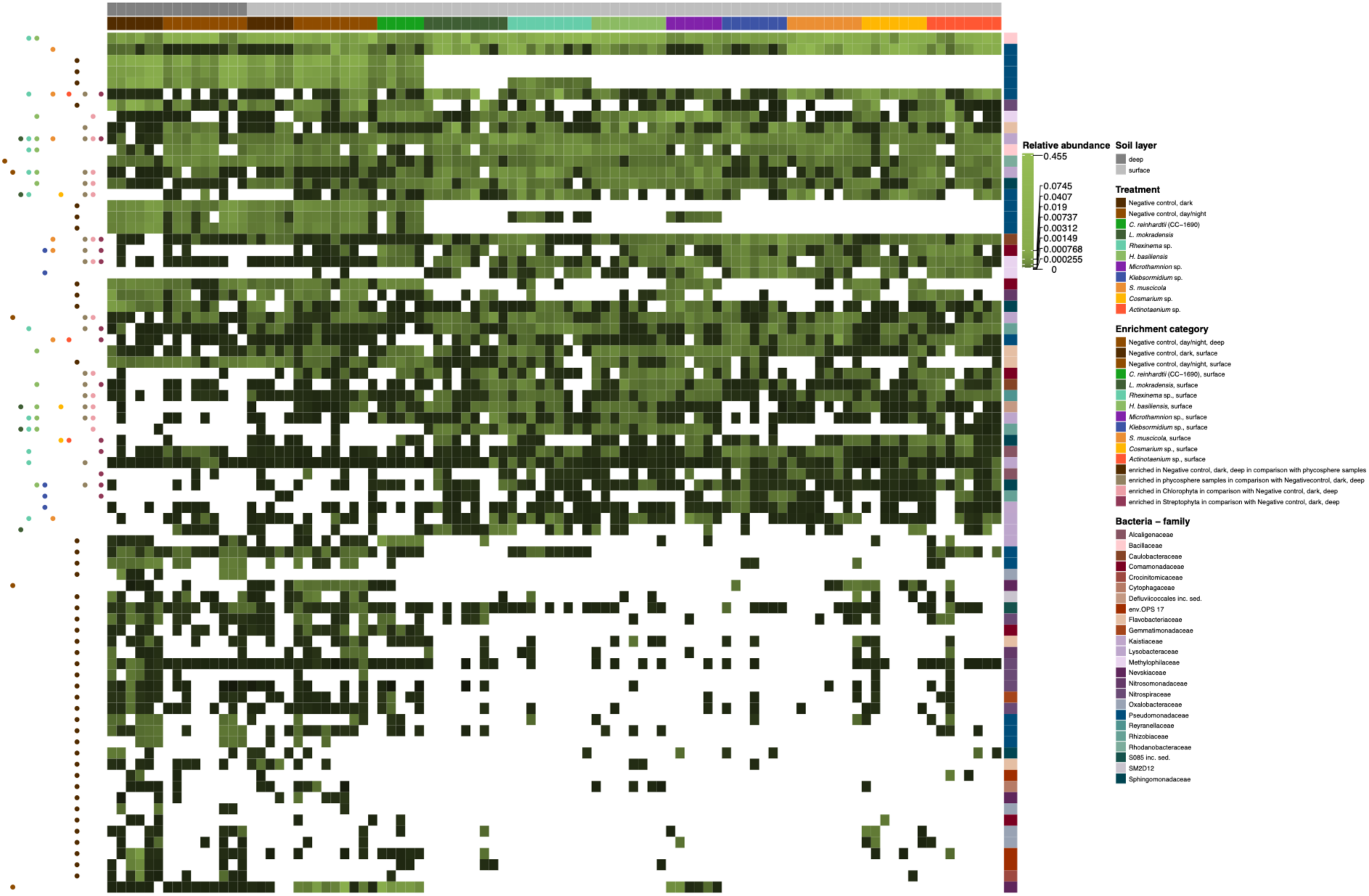
Heatmap of relative abundances of ASVs enriched in negative controls, all phycospheres, Streptophyta or Chlorophyta phycospheres and single alga phycospheres. Left annotation indicates the enriched compartments, and right annotation - the bacterial family the ASV belongs to.

**Figure S9.**
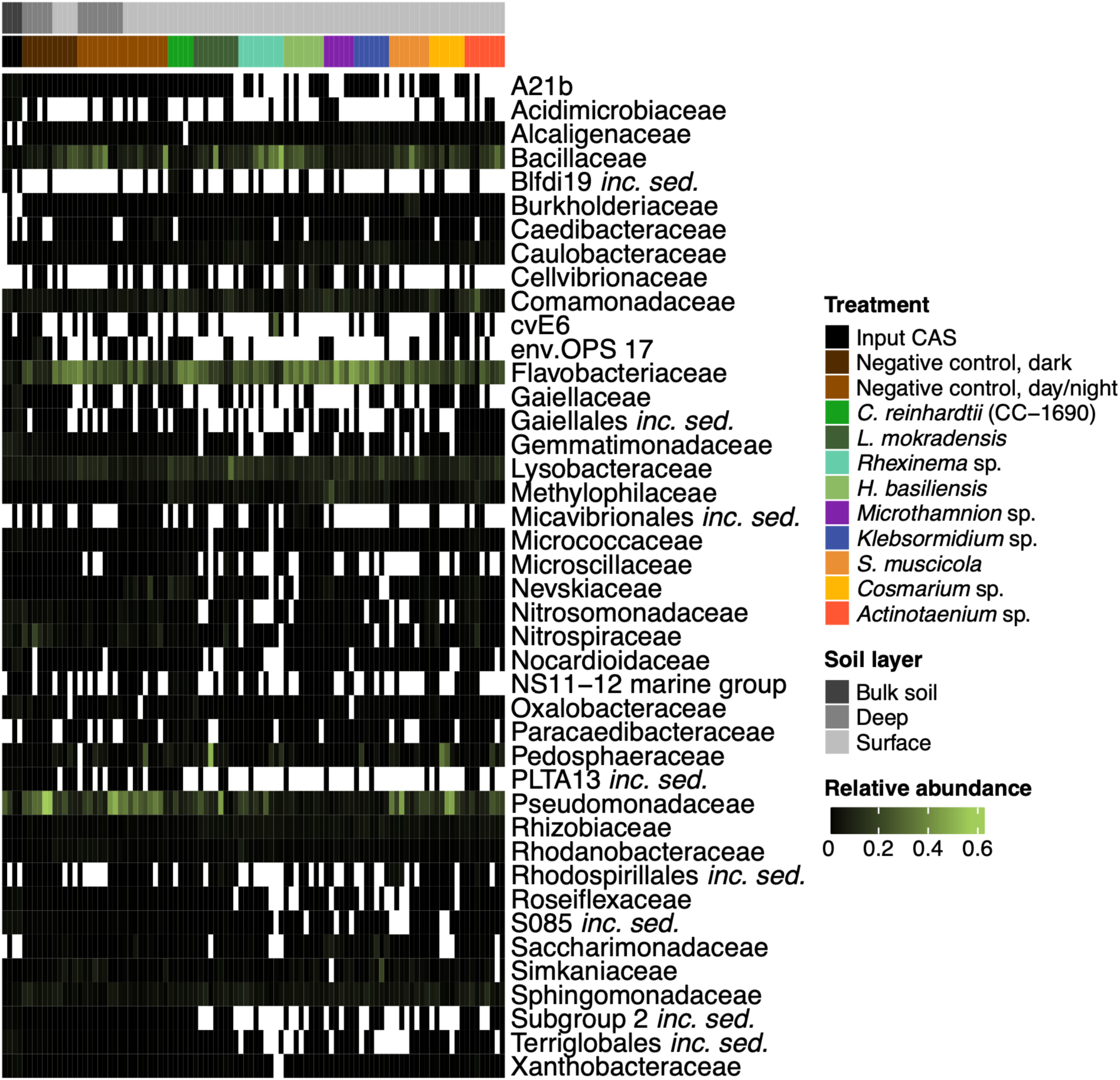
Heatmap of bacterial family-level relative abundances in input CAS, negative controls and phycosphere samples. The heatmap shows relative abundances of bacterial families (rows) across samples (columns). Only families with ≥1% mean relative abundance in at least one condition are included. Columns are annotated by Treatment and Soil layer (top annotation). Color intensity indicates relative abundance, with white representing zero abundance (family not detected in the sample).

**Figure S10.**
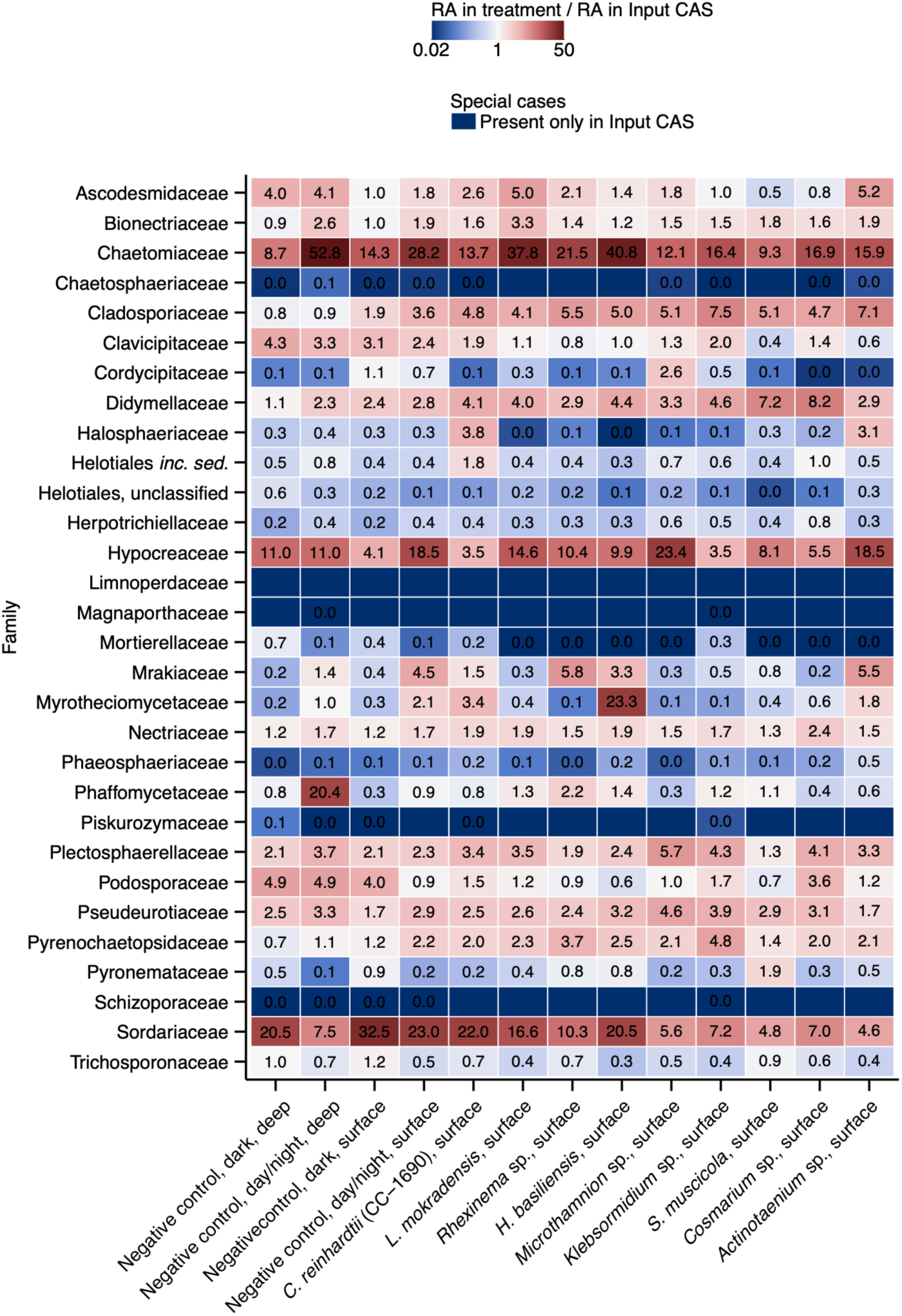
Ratios of mean family-level fungal RA between phycospheres or negative controls and input CAS (RA_treatment_/RA_input_). Cells are colored according to the ratio: shades of blue indicate a decrease in family-level RA in the phycosphere, and shades of red indicate an increase. Numbers represent the ratio value; families detected only in the phycospheres or input CAS are shown without a number. Only families with ≥1% mean relative abundance in either input CAS or one of the treatments are included.

**Figure S11.**
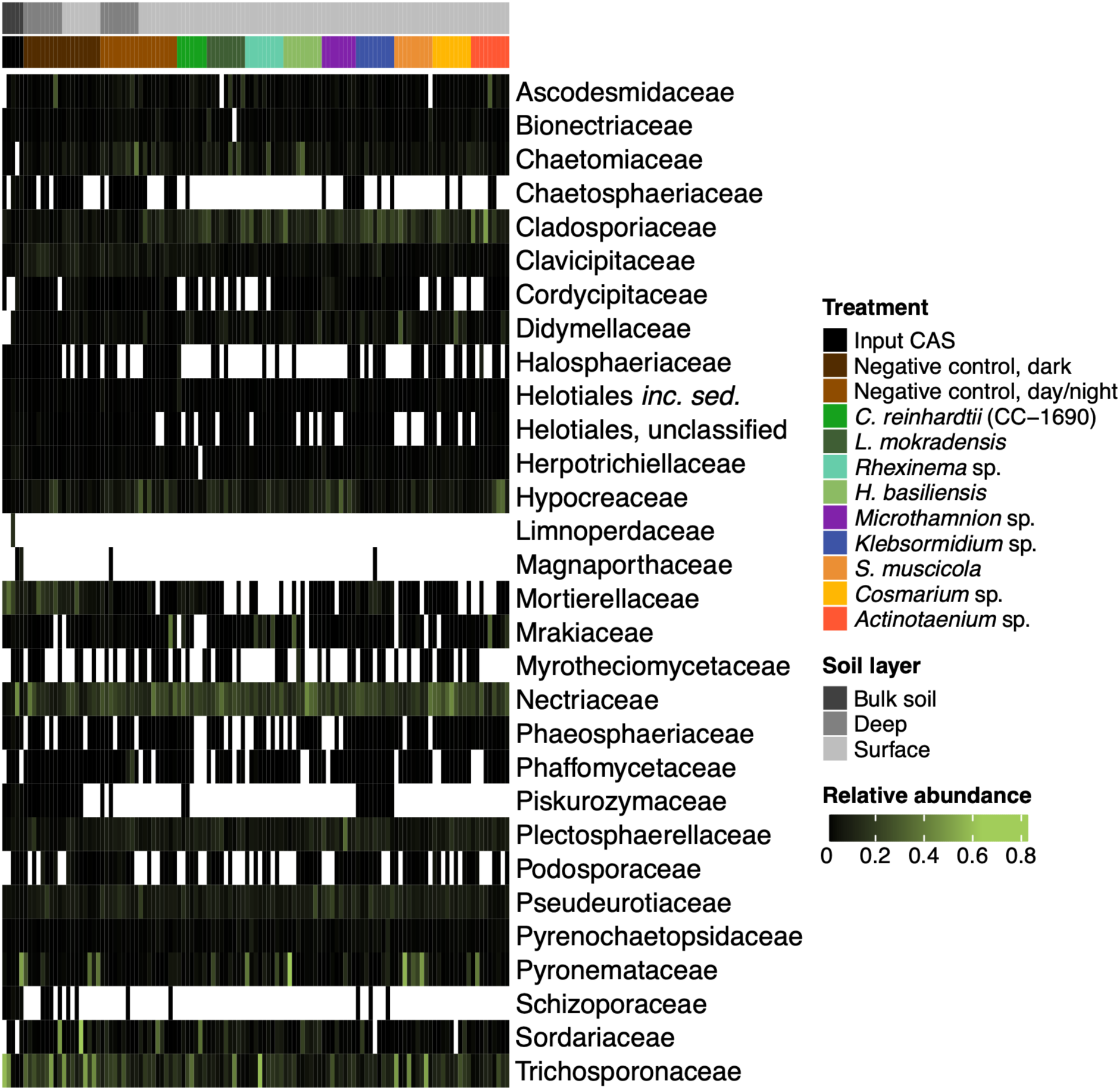
Heatmap of fungal family-level relative abundances in input CAS, negative controls and phycosphere samples. The heatmap shows relative abundances of fungal families (rows) across samples (columns). Only families with ≥1% mean relative abundance in at least one condition are included. Columns are annotated by Treatment (top). Color intensity indicates relative abundance, with white representing zero abundance (family not detected in the sample).

**Figure S12.**
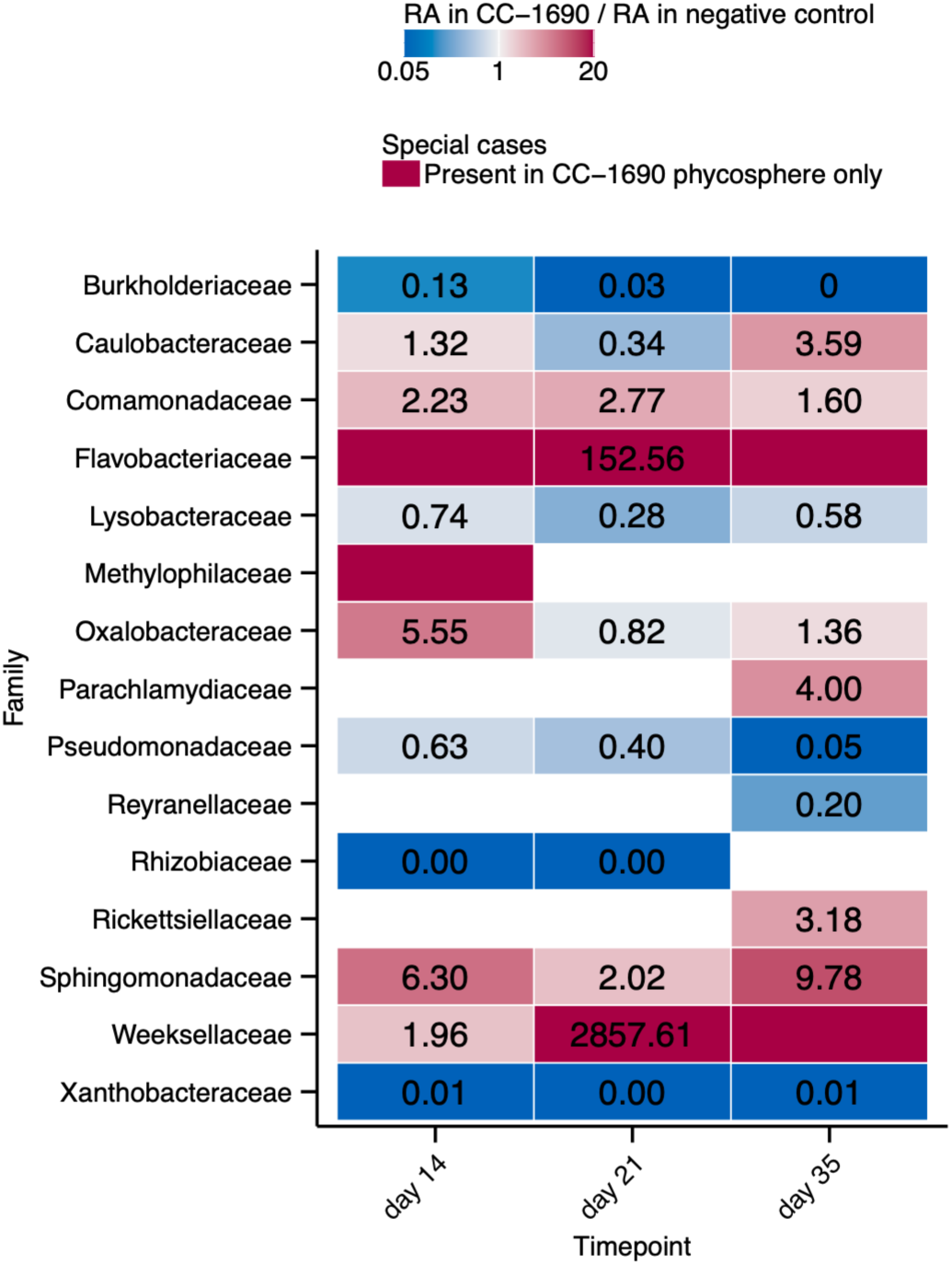
Ratios of mean family-level bacterial RA between *C. reinhardtii* CC-1690 phycosphere and negative control at different timepoints (RA_treatment_/RA_control_). Cells are colored according to the ratio: shades of blue indicate a decrease in family-level RA in the phycosphere, and shades of red indicate an increase. Numbers represent the ratio value; families detected only in the phycosphere are shown without a number. White cells indicate families absent from both the control and phycosphere at a given time point. Only families with ≥1% mean relative abundance in either control or *C. reinhardtii* CC-1690 phycosphere are included.

**Figure S13.**
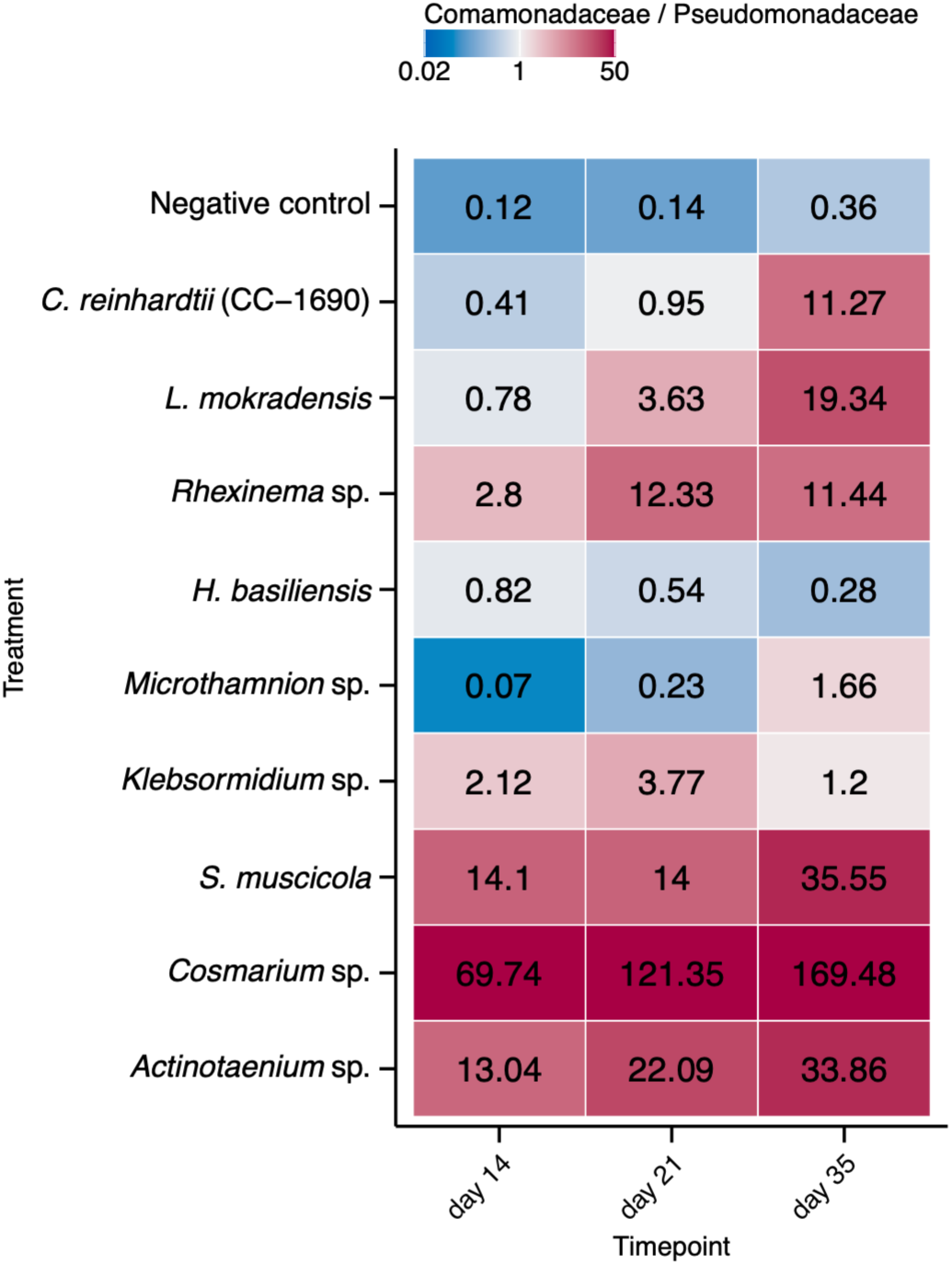
Ratios of mean Comamonadaceae RA to *Pseudomonadaceae* RA for different treatments (RA_Comamonadaceae_/RA_Pseudomonadaceae_). Cells are colored according to the ratio: shades of blue indicate Pseudomonadaceae domination, and shades of red indicate *Comamonadaceae* domination. Numbers represent the ratio value.

**Figure S14.**
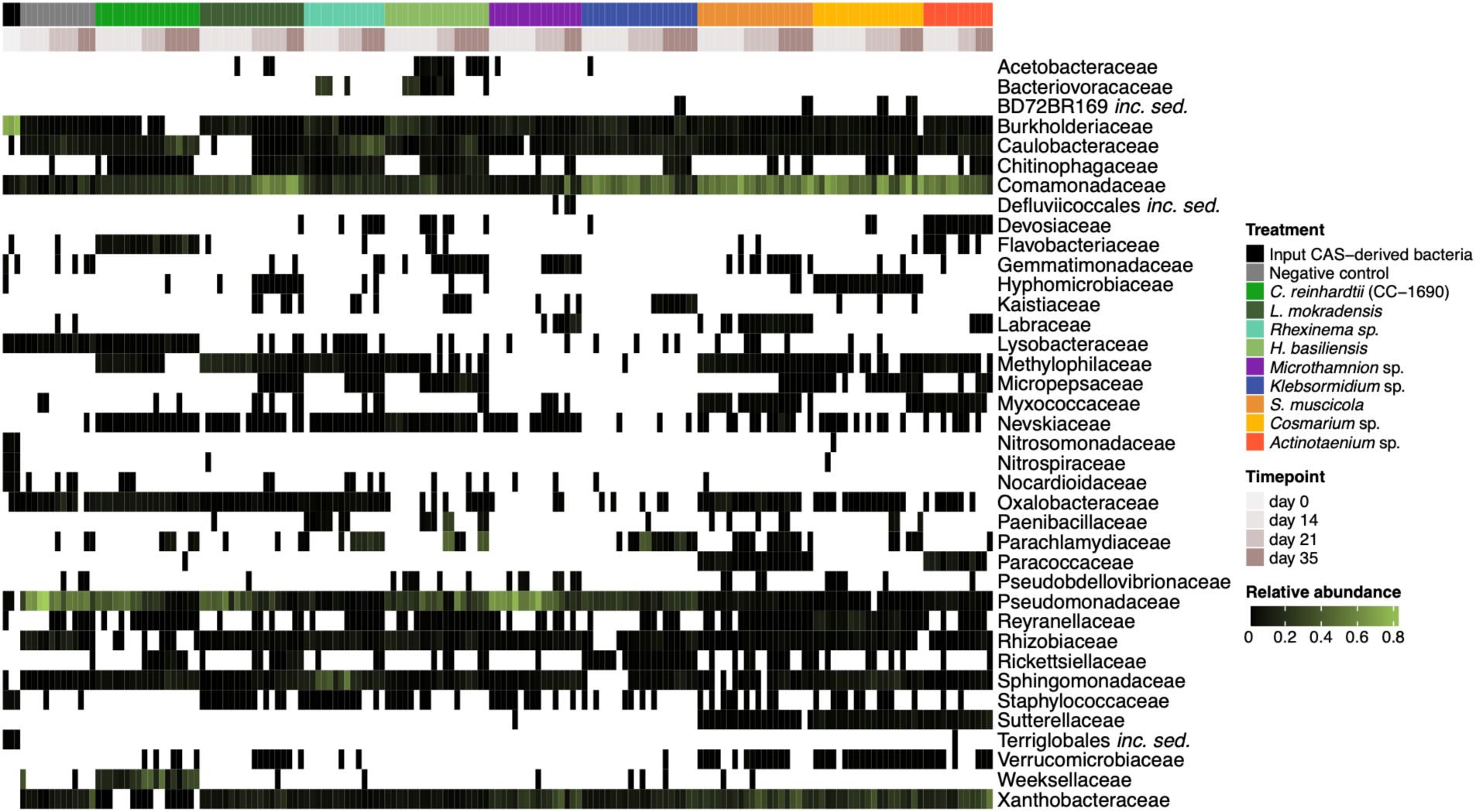
Heatmap of bacterial family-level relative abundances in input CAS-derived bacteria, negative controls and phycosphere samples. The heatmap shows relative abundances of bacterial families (rows) across samples (columns). Only families with ≥1% mean relative abundance in at least one condition are included. Columns are annotated by Treatment (top). Color intensity indicates relative abundance, with white representing zero abundance (family not detected in the sample).

**Figure S15.**
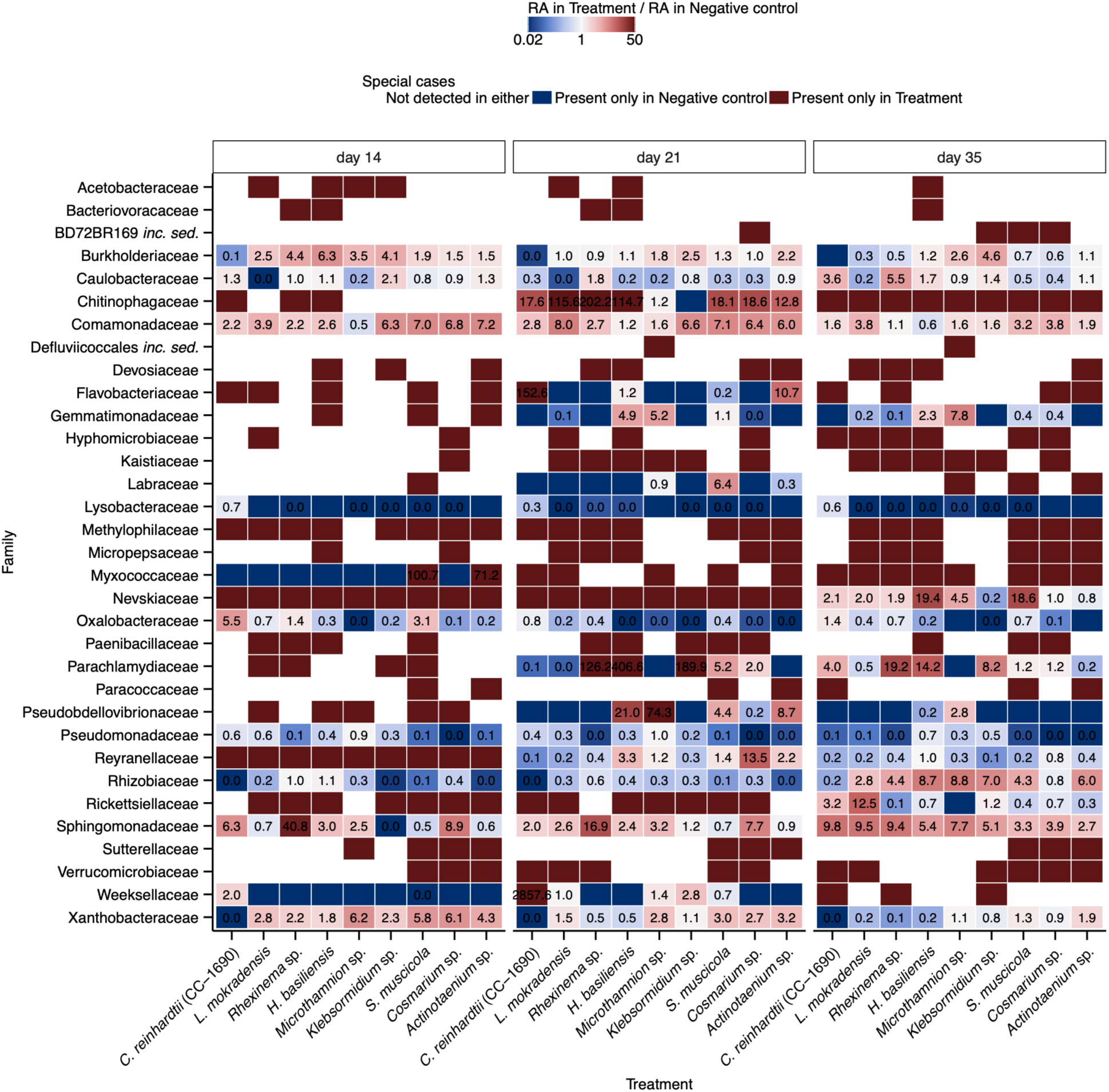
Ratios of mean family-level bacterial RA between phycospheres and negative control at a given timepoint (RA_treatment_/RA_control_). Cells are colored according to the ratio for a given family: shades of blue indicate a lower RA in the phycosphere, and shades of red indicate a higher RA in the phycosphere. Numbers represent the ratio value; families detected only in the phycospheres or controls are shown without a number. White cells indicate families absent from both the control and phycosphere at a given time point. Only families with ≥1% mean relative abundance in either control or one of the treatments are included.

**Figure S16.**
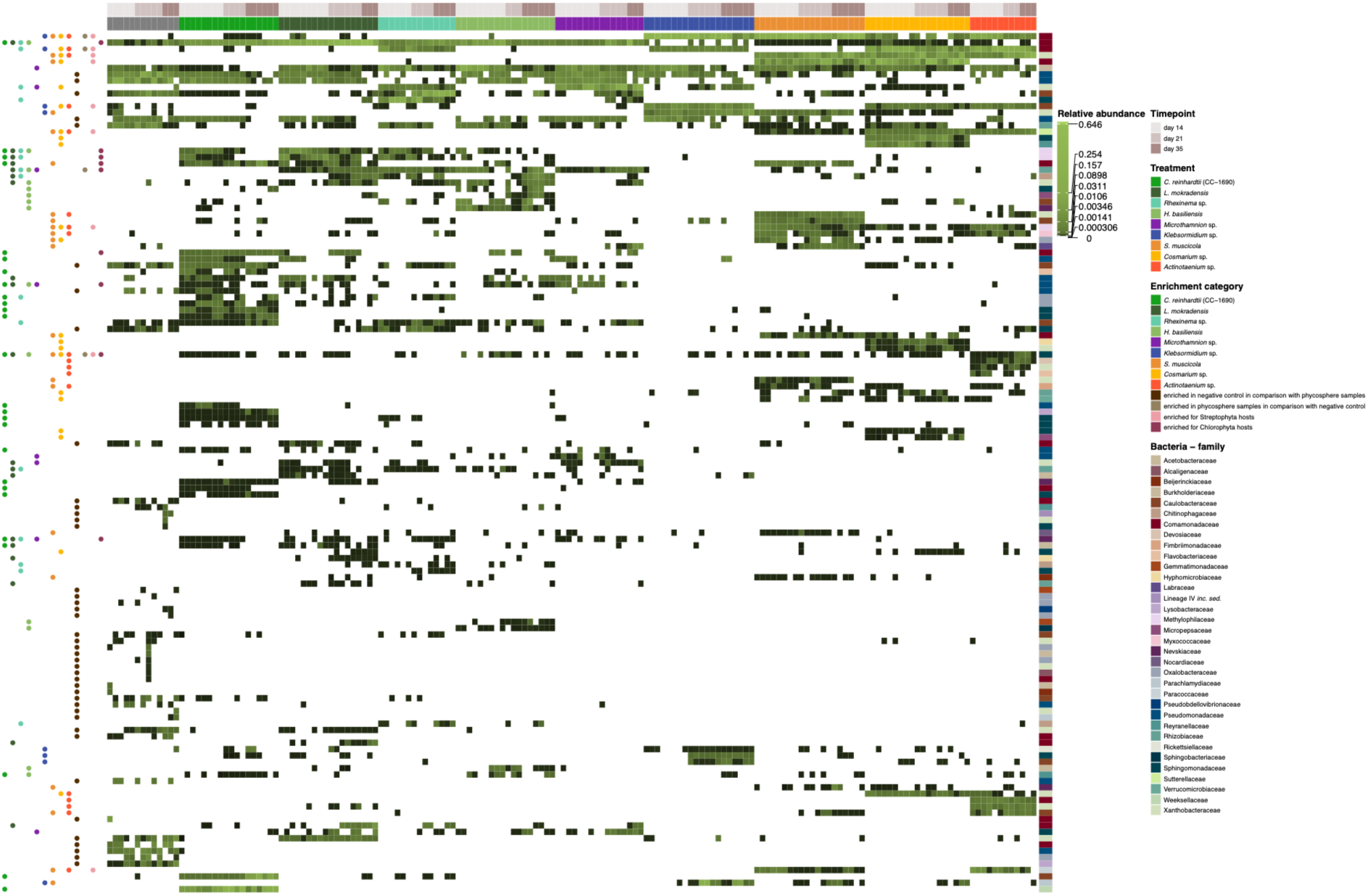
Heatmap showing relative abundances of ASVs enriched in negative controls, all phycospheres, Streptophyta or Chlorophyta phycospheres and single alga phycospheres. Left annotation indicates the enriched compartments, and right annotation - the bacterial family the ASV belongs to.

## Supplementary tables

**Supplementary Table 1.** Algae used in this study.

**Supplementary Table 2.** Mantel test of correlations between microbial Bray-Curtis dissimilarities and algal phylogenetic distances.

**Supplementary Table 3.** Primers used in this study. Supplementary Table 4. PCR programs.

**Supplementary Table 5.** Initial (morphology based) classification of the algal isolates and current classification base don algal morphology and molecular identification.

**Supplementary Table 6. Composition of buffers used for SPRI beads DNA extraction.**

## Notes

### Competing Interest Statement

The authors have declared no competing interest.

## References

Abarenkov, K. et al. The UNITE database for molecular identification and taxonomic communication of fungi and other eukaryotes: sequences, taxa and classifications reconsidered. Nucleic Acids Res. 52, D791–D797 (2024).

Amin, S. A. et al. Photolysis of iron-siderophore chelates promotes bacterial-algal mutualism. Proc. Natl. Acad. Sci. U. S. A. 106, 17071–17076 (2009).

Amin, S. A. et al. Interaction and signalling between a cosmopolitan phytoplankton and associated bacteria. Nature 522, 98–101 (2015).

Bai, Y. et al. Functional overlap of the Arabidopsis leaf and root microbiota. Nature 528, 364–369 (2015).

Bäcker, M, et al. Spatial Structure: Shaping the Ecology and Evolution of Microbial Communities. FEMS Microbiology Reviews 50 fuaf067 (2026)

Beckers, B. et al. Performance of 16s rDNA primer pairs in the study of rhizosphere and endosphere bacterial microbiomes in metabarcoding studies. Front. Microbiol. 7, 650 (2016).

Bolyen, E. et al. Reproducible, interactive, scalable and extensible microbiome data science using QIIME 2. Nat. Biotechnol. 37, 852–857 (2019).

Bonito, G. et al. Plant host and soil origin influence fungal and bacterial assemblages in the roots of woody plants. Mol. Ecol. 23, 3356–3370 (2014).

Bourceret, A. et al. Maize field study reveals covaried Microbiota and metabolic changes in roots over plant growth. MBio 13, e0258421 (2022).

Caesar, J., Tamm, A., Ruckteschler, N., Leifke, A. L. & Weber, B. Revisiting chlorophyll extraction methods in biological soil crusts -methodology for determination of chlorophyll *a* and chlorophyll *a* + *b* as compared to previous methods. Biogeosciences 15, 1415– 1424 (2018).

Calatrava, V. et al. Genetic evidence for algal auxin production in Chlamydomonas and its role in algal-bacterial mutualism. iScience 27, 108762 (2024).

Callahan, B. J. et al. DADA2: High-resolution sample inference from Illumina amplicon data. Nat. Methods 13, 581–583 (2016).

Carrión, V. J. et al. Pathogen-induced activation of disease-suppressive functions in the endophytic root microbiome. Science 366, 606–612 (2019).

Castrillo, G. et al. Root microbiota drive direct integration of phosphate stress and immunity. Nature 543, 513–518 (2017).

Chelius, M. K. & Triplett, E. W. The Diversity of Archaea and Bacteria in Association with the Roots of Zea mays L. Microb Ecol 41, 252–263 (2001).

Chen, Y. et al. Soil fungal communities show more specificity than bacteria for plant species composition in a temperate forest in China. BMC Microbiol. 22, 208 (2022).

Cheng, S. et al. Genomes of Subaerial Zygnematophyceae Provide Insights into Land Plant Evolution. Cell 179, 1057–1067.e14 (2019).

Croft, M. T., Lawrence, A. D., Raux-Deery, E., Warren, M. J. & Smith, A. G. Algae acquire vitamin B12 through a symbiotic relationship with bacteria. Nature 438, 90–93 (2005).

Delaux, P.-M. et al. Algal ancestor of land plants was preadapted for symbiosis. Proc. Natl. Acad. Sci. U. S. A. 112, 13390–13395 (2015).

Delaux, P.-M. & Schornack, S. Plant evolution driven by interactions with symbiotic and pathogenic microbes. Science 371, eaba6605 (2021).

Durham, B. P. et al. Cryptic carbon and sulfur cycling between surface ocean plankton. Proc. Natl. Acad. Sci. U. S. A. 112, 453–457 (2015).

Durán, P. et al. Microbial Interkingdom Interactions in Roots Promote Arabidopsis Survival. Cell 175, 973–983.e14 (2018).

Durán, P. et al. Shared features and reciprocal complementation of the Chlamydomonas and Arabidopsis microbiota. Nat. Commun. 13, 1–14 (2022).

Durán, P. The core microbiota across the green lineage. Curr. Opin. Plant Biol. 77, 102487 (2024).

Fay, M. F., Swensen, S. M. & Chase, M. W. Taxonomic Affinities of Medusagyne oppositifolia (Medusagynaceae). Kew Bull. 52, 111 (1997).

Fei, C. et al. Quorum sensing regulates “swim-or-stick” lifestyle in the phycosphere. Environ. Microbiol. 22, 4761–4778 (2020).

Fitzpatrick, C. R. et al. Assembly and ecological function of the root microbiome across angiosperm plant species. Proc. Natl. Acad. Sci. U. S. A. 115, E1157–E1165 (2018).

Flores, D. C. et al. A mutualistic bacterium rescues a green alga from an antagonist. Proceedings of the National Academy of Sciences 121, e2401632121 (2024).

Friedlingstein, P. et al. Global carbon budget 2019. Earth Syst. Sci. Data 11, 1783–1838 (2019).

Gontcharov, A. A., Marin, B. & Melkonian, M. Are combined analyzes better than single gene phylogenies? A case study using SSU rDNA and rbcL sequence comparisons in the Zygnematophyceae (Streptophyta). Mol. Biol. Evol. 21, 612–624 (2004).

Fu, He, Mario Uchimiya, Jeff Gore, and Mary Ann Moran. 2020. Ecological Drivers of Bacterial Community Assembly in Synthetic Phycospheres. Proceedings of the National Academy of Sciences of the United States of America 117 (7): 3656–62.

Grafen, A. The phylogenetic regression. Philos. Trans. R. Soc. Lond. B Biol. Sci. 326, 119–157 (1989).

Garrigues, V., Guan, R., Preteseille, N., Leccia, I., Wagner, E., Roux, F., Duran, P. Environmental factors and microbe-microbe interactions drive the structure of the core microbiota of terrestrial microalgae. BioRxiv, 2026-09.

Goldford, Joshua E., Nanxi Lu, Djordje Bajić, Sylvie Estrela, Mikhail Tikhonov, Alicia Sanchez-Gorostiaga, Daniel Segrè, Pankaj Mehta, and Alvaro Sanchez. 2018. “Emergent Simplicity in Microbial Community Assembly.” Science 361 (6401): 469–74.

Harbort, C. J. et al. Root-Secreted Coumarins and the Microbiota Interact to Improve Iron Nutrition in Arabidopsis. Cell Host Microbe 28, 825–837.e6 (2020).

Jassey, V. E. J. et al. Contribution of soil algae to the global carbon cycle. New Phytol. 234, 64–76 (2022).

Kim, H. et al. Bacterial response to spatial gradients of algal-derived nutrients in a porous microplate. ISME J. 16, 1036–1045 (2022).

Kimbrel, J. A. et al. Host selection and stochastic effects influence bacterial community assembly on the microalgal phycosphere. Algal Res. 40, 101489 (2019).

Knack, J. J. et al. Microbiomes of streptophyte algae and bryophytes suggest that a functional suite of Microbiota fostered plant colonization of land. Int. J. Plant Sci. 176, 405–420 (2015).

Manhart, J. R. Phylogenetic analysis of green plant rbcL sequences. Mol. Phylogenet. Evol. 3, 114–127 (1994).

Marin, B., Klingberg, M. & Melkonian, M. Phylogenetic relationships among the Cryptophyta: Analyzes of nuclear-encoded SSU rRNA sequences support the monophyly of extant Plastid-containing lineages. Protist 149, 265–276 (1998).

Martin, M. Cutadapt removes adapter sequences from high-throughput sequencing reads. EMBnet J. 17, 10 (2011).

Mazel, F. et al. Is host filtering the main driver of phylosymbiosis across the tree of life? mSystems 3, e00097–18 (2018).

McCourt, R. M. et al. PHYLOGENY OF THE CONJUGATING GREEN ALGAE (ZYGNEMOPHYCEAE) BASED ON rbc L SEQUENCES. J. Phycol. 36, 747–758 (2000).

Medlin, L., Elwood, H. J., Stickel, S. & Sogin, M. L. The characterization of enzymatically amplified eukaryotic 16S-like rRNA-coding regions. Gene 71, 491–499 (1988).

Barbara, M. & Michael, M. Three new synthetic algal culture media to grow them all. Biobreeding 1, 10005–10005 (2026).

Meyberg, M. 1988. “Selective Staining of Fungal Hyphae in Parasitic and Symbiotic Plant-Fungus Associations.” Histochemistry 88 (**2**): 197–99.

Naylor, D., DeGraaf, S., Purdom, E. & Coleman-Derr, D. Drought and host selection influence bacterial community dynamics in the grass root microbiome. ISME J. 11, 2691–2704 (2017).

Okonechnikov, K., Golosova, O., Fursov, M. & UGENE team. Unipro UGENE: a unified bioinformatics toolkit. Bioinformatics 28, 1166–1167 (2012).

Community Ecology Package [R package vegan version 2.7-2]. Comprehensive R Archive Network (CRAN) https://CRAN.R-project.org/package=vegan (2025).

Paerl, R. W. et al. Use of plankton-derived vitamin B1 precursors, especially thiazole-related precursor, by key marine picoeukaryotic phytoplankton. ISME J. 11, 753–765 (2016).

Paradis, E. & Schliep, K. ape 5.0: an environment for modern phylogenetics and evolutionary analyzes in R. Bioinformatics 35, 526–528 (2019).

Pershina, E. et al. Comparative analysis of prokaryotic communities associated with organic and conventional farming systems. PLoS One 10, e0145072 (2015).

Quast, C. et al. The SILVA ribosomal RNA gene database project: improved data processing and web-based tools. Nucleic Acids Res. 41, D590–6 (2013).

R Core Team. R: A Language and Environment for Statistical Computing. (R Foundation for Statistical Computing, Vienna, Austria, 2024).

Robeson, M. S 2nd., et al. RESCRIPt: Reproducible sequence taxonomy reference database management. PLoS Comput. Biol. 17, e1009581 (2021).

Sasso, S., et al. From molecular manipulation of domesticated Chlamydomonas reinhardtii to survival in nature. Elife 7, (2018).

Seymour, J. R., et al. Zooming in on the phycosphere: the ecological interface for phytoplankton-bacteria relationships. Nat Microbiol 2, 17065 (2017).

Shen, W. et al. SeqKit: A cross-platform and ultrafast toolkit for FASTA/Q file manipulation. PLoS One 11, e0163962 (2016).

Simmons, T. et al. Drought Drives Spatial Variation in the Millet Root Microbiome. Front. Plant Sci. 11, 599 (2020).

Stock, W., Willems, A., Mangelinckx, S., Vyverman, W. & Sabbe, K. Selection constrains lottery assembly in the microbiomes of closely related diatom species. ISME Commun. 2, 11 (2022).

Thiergart, T. et al. Root microbiota assembly and adaptive differentiation among European Arabidopsis populations. *Nat*. Ecol. Evol. 4, 122–131 (2020).

Tkacz, A. et al. Influence of plant fraction, soil, and plant species on Microbiota: A multikingdom comparison. MBio 11, (2020).

Wang, Chaoqun, and Yakov Kuzyakov. 2024. Mechanisms and Implications of Bacterial-Fungal Competition for Soil Resources. The ISME Journal 18 (1): wrae073.

Wippel, K. et al. Host preference and invasiveness of commensal bacteria in the Lotus and Arabidopsis root microbiota. Nat Microbiol 6, 1150–1162 (2021).

Xu, L. et al. Drought delays development of the sorghum root microbiome and enriches for monoderm bacteria. Proceedings of the National Academy of Sciences 115, E4284– E4293 (2018).

Yilmaz, P. et al. The SILVA and “All-species Living Tree Project (LTP)” taxonomic frameworks. Nucleic Acids Res. 42, D643–8 (2014).

